# Systematic Characterization of Transport Regulation in *Escherichia coli* across defined environmental cues

**DOI:** 10.1101/2024.08.26.609649

**Authors:** Christoph Binsfeld, Roberto Olayo Alarcon, Morgane Wartel, Mara Stadler, Christian Müller, Ana Rita Brochado

## Abstract

Transport of small molecules across the bacterial cell envelope is essential to ensure nutrient uptake and protect bacteria from toxic compounds. Due to its decisive role, transport is controlled by complex regulatory networks, knowledge of which – in particular across external cues – remains poorly understood. Here we investigate transcriptional regulation of seven prominent transport genes in *Escherichia coli* across 94 defined chemical cues, and simultaneously map the contributions of the key regulators MarA, SoxS and Rob to promoter activity. One third of all tested compounds triggered transcriptional changes, the vast majority of which previously unknown. Importantly, we exposed main drivers of transport control in *E. coli*, e.g. bacteriostatic, but not bactericidal, antibiotics trigger expression of efflux pumps, and that Rob contributes to ∼1/3 of all measured transcriptional changes, thereby emerging as a more prominent transport regulator than previously thought. We showcase the potential of our resource by elucidating the molecular mechanism of antibiotic antagonisms with widely consumed caffeine in *E. coli*. Altogether, our resource capitalizes on providing a quantitative overview of transport determinants across environments, and brings perspective to long-term prevailing concepts in the field.

## Introduction

Gram-negative bacteria have a double membrane surrounding their cell wall, which acts as a selective barrier against the environment. Influx and efflux of chemically diverse small molecules, including harmful substances, but also nutrients and intracellular metabolites, across the cell envelope is facilitated by protein-channels, namely outer membrane porins (OMPs) and efflux pumps^1,2^. A delicate balance between porin-mediated passive uptake and a fast active efflux imposes a permeability barrier, and it is therefore a crucial determinant for bacteria to thrive in harsh environments. Numerous mutations in efflux pumps, porins and their regulatory elements have been associated with antibiotic resistance in clinical isolates^2–5^. Furthermore, pump deletion mutants have impaired host-intracellular survival^6^ and restricted antibiotic persistence phenotypes^7–9^. It is also becoming increasingly clear that the same import/export machineries are determinant for sensitivity to non-antibiotic drugs, not only in pathogenic bacteria but also in commensal members of bacterial communities, such as the gut microbiota^10–12^. Our own previous work suggests that transport regulation is a critical determinant of synergy and antagonism (drug interactions) in bacteria, at least as important as the antibiotic targets themselves^13,14^. Yet, our knowledge on how bacteria control their import/export machineries across environments remains limited, preventing better design of treatment strategies.

Three major proteins, OmpC, OmpF and PhoE, represent a substantial fraction of the total number of outer membrane porins in *E. coli*. Together, they import a wide range of anionic (PhoE) and cationic small molecules (OmpC, OmpF), including clinically important antibiotics, such as β-lactams and fluroquinolones^5^. In what efflux is concerned, six efflux pump families have been described in bacteria, using either ATP or electrochemical gradients as energy source for active transport. The resistance-nodulation-cell division (RND) pump AcrAB-TolC is among the most well-characterized efflux pumps in *Enterobacteriaceae*, including *E. coli*^2^. This is a tri-partite protein complex spanning the inner- and outer-membrane effluxing a wide range of antibiotics, such as β-lactams, fluroquinolones, macrolides among others^2^. Due to their fundamental role in controlling membrane permeability, influx and efflux machineries are highly regulated at transcriptional and post-transcriptional levels^2^. Regulatory mechanisms include two-component systems (e.g. CpxAR), small proteins that modulate pump specificity (e.g. AcrZ), and global transcriptional regulators such as the transcription factors MarA, SoxS, Rob and RamA (latter absent in *E. coli*). MarA, SoxS and Rob bind the so called mar-sox-rob box, a degenerate sequence of ∼20 base-pairs found in multiple promoter sequences of over 40 genes in *E. coli*, including *acrAB, tolC* and *micF,* which encodes *a* small RNA involved in porin regulation (Fig. 1a)^15^. This arrangement enables *E. coli* to orchestrate a complex transcriptional network in response to different environmental cues, such as oxidative stress, toxic compounds, acidic pH, bile acids and more^2,15–18^. Extensive efforts have been directed to structurally and functionally characterize several components of this network, including transcriptome analysis^19–22^, regulator binding sites using chromatin immunoprecipitation and DNA sequencing^23^, identification of canonical chemical cues^16–18,24,25^, ligand-regulator structure determination^26,27^, pumps and porins specificity studies^28,29^, among many others. Efflux pump inhibitors, ideal compounds for potentiating antibiotic activity by directly impairing efflux, have been identified^30^. However, given the complexity, redundancy and extensive cross-talk inherent to this regulatory network^2,15^ (Fig. 1a), current knowledge is still not sufficient to enable full understanding of network operational modes, namely whether and how the hierarchy of regulator-responsive promoters changes across environmental cues.

**Figure 1:**
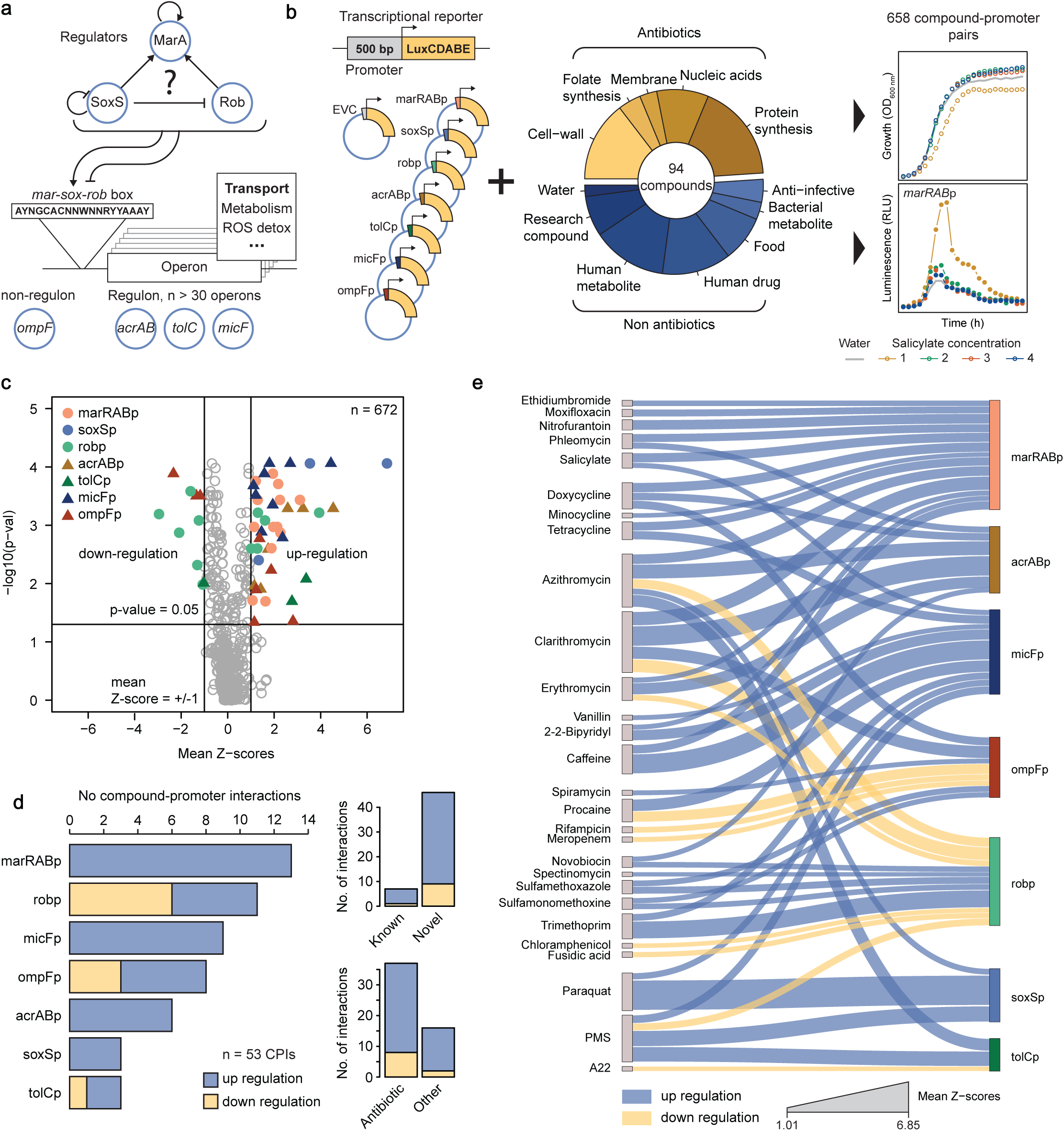
Unravelling transcriptional regulation of drug transport-related genes in E. coli under chemical stress. **a)** Simplified schematic representation of the Mar-Sox-Rob network, as well as placement of the 7 promoters selected for this study. Previously described auto- and cross-regulation between the three regulators^2,37^, as well as their degenerate binding sequence (mar-sox-rob box^52^) are also represented. **b)** Schematic overview of screening approach. Lux-based transcriptional reporters of 7 key transport-related genes and the compound library used in this study to probe 658 compound-promoter interactions. Growth and luminescence were periodically measured over 12 h. **c)** Compound-promoter interactions in *E. coli*. Volcano plot summarizing the screen results shows 53 significant CPIs (colored by promoter) amongst 658 tested (+water, n=672 in total). X-axis: mean Z-scores (n=4 concentrations x 2 biological replicates = 8). Y-axis: Benjamini Hochberg adjusted p-value of double-sided rank-sum statistical tests between Z-scores of compound-promoter pairs (n=8) and water (n=16). **d)** General features of compound-promoter interactions (CPIs). Number (No) of interactions per promoter (left), their classification according to novelty (upper right corner) and whether they involve an antibiotic (lower right corner) are shown. **e)** CPI network. 53 significant compound- promoter interactions are shown as edges in a Sankey diagram connecting the compounds (left, source nodes) to the promoters (right, target nodes). Edge thickness represents mean Z-scores (n = 8), while node size represents the total number of interactions.

Here we propose a systematic and quantitative approach to elucidate transcriptional control of 7 key transport genes in the model organism *E. coli* across 94 chemically defined environments. Our results expose several new modulators of bacterial transport, such as the macrolide antibiotics, but also non-antibiotic compounds such as caffeine. In a complementary experimental approach using regulator deletion mutants, we quantified regulator contribution to promoter activity under all 94 compounds using a simple regression model, and mapped how the same regulators can promote or hinder transcriptional regulation of a target gene depending on the chemical cue. We capitalize on our unique systematic approach to uncover general drivers of transport regulation in *E. coli*, for instance that Rob has a much more prevalent role in transcriptional control of transport than previously acknowledged. Our findings illustrate how a comprehensive approach is crucial for revealing operational modes of highly complex regulatory networks, such as the one at hand. Finally, we showcase the mechanistic potential of our approach by elucidating the molecular mechanism by which caffeine, a widely consumed food ingredient, induces low level resistance to fluroquinolones via decreased uptake.

## Results

### Unravelling transcriptional regulation of transport-related genes in *E. coli* under chemical stress

We set out to systematically investigate the transcriptional response of the most prominent genes controlling transport in *E. coli* across 94 defined chemical stresses in a concentration resolved manner. We constructed plasmid-based luminescence reporters for 7 genes, 6 of which contain the degenerate mar-sox-rob box in their promoter region (Fig. 1a): the key transcription factors *marA*, *soxS* and *rob*, *acrAB* and *tolC*, which are both components of the major efflux pump AcrAB-TolC, and the small-RNA *micF*, a post-transcriptional regulator of *ompF*^31^. In addition, we included a reporter of *ompF* transcription, to gain insight on how vast is the transcriptional adjustment of transport related genes which are not under direct control of *marA*, *soxS* or *rob* (Methods, supplementary table 1). In total, we probed 658 compound-promoter pairs. As many compounds, and specifically antibiotics, are known to cause vast transcriptional effects^32,33^, we included a reporter strain containing a promoterless/leaky luminescence reporter (empty vector control, EVC) to better control for possible non-specific transcriptional effects of each compound. We assembled a compound collection containing 94 compounds including all major antibiotic classes, human-targeted drugs (e.g aspirin), gut metabolites (e.g. bile acids) and small-molecules found in common foods (e.g. vanillin, Fig. 1b, supplementary table 2). We ensured high overlap of the compound library with our previous work on systematic assessment of drug combinations in Gram-negative bacteria (61 out of 79 compounds)^13^, to enable downstream integration of the datasets for interpretation purposes. All compound-promoter pairs were probed in duplicates across four compound concentrations (2-fold dilutions, ED. Fig. 1a). Maximum concentrations were adjusted to be close to minimum inhibitory concentration (MIC) for antimicrobials, 500 µM for most non-antimicrobials, and up to 1 mM for small compounds with similarity to canonical inducers (positive controls, e.g. salicylate, supplementary table 2, Methods). Briefly, growth (optical density, OD_600 nm_) and luminescence for each reporter strain were periodically monitored over 12 hours in the presence of each individual compound at all concentrations (Methods). Area under the curve for a period of 8h (AUC) was used as a proxy for growth (OD_AUC_) and luminescence (Lux_AUC_) profiles (onset of stationary phase, ED. Fig. 1b). As expected, growth was reasonably constant across all reporter strains, while luminescence showed a large dynamic range depending on the promoter, with tolCp showing the lowest and ompFp the highest signal (ED. Fig. 1c&d). We excluded the possibility that some compounds, namely protein synthesis inhibitors, could increase plasmid copy number (as previously reported for different origin of replication^34^) by quantitative PCR (ED. Fig. 1e). High data quality is reflected by Pearson correlation between replicates above 0.8 across all promoters, except for tolCp, likely due to its weak promoter strength (ED. Fig. 1f). Nonetheless, since the effect of compounds expected to trigger its expression via oxidative stress – e.g. phenazine methosulfate – could be captured here, we decided to keep tolCp in our dataset.

Next we developed a data analysis pipeline to systematically assess *compound-promoter* interactions, which we define as increased or decreased *promoter* activity as measured by our luminescence reporter in the presence of the *compound* (Methods, ED. Fig. 1b). Briefly, we computed an *interaction score* for any given *compound-promoter* pair based on its deviation of normalized luminescence (Lux_AUC_/OD_AUC_) to the c*ompound-EVC*. We subsequently Z-transformed the interaction scores (Z-scores) to allow comparability of promoters of varying signal intensity. Finally, significant *Compound-Promoter Interactions* (CPIs) were called based on a double cutoff on mean Z-scores and rank-sum test p-value comparing the Z-score distributions of each compound-promoter with that of water-promoter (Fig. 1c, Methods). We identified 53 CPIs distributed across all promoters and 28 out of the 94 compounds (ED Fig. 2a), with induction being stronger and more prevalent than repression (43 vs 10 instances, Fig. 1d). While the majority of the 28 compounds are antibiotics, ∼1/3 of all identified CPIs involve non-antibiotic compounds, showing that non-antibiotics also modulate transport across the cell envelope^12,13^ (Fig 1d). The number of interactions per promoter varies between three for soxSp and tolCp and 13 for marRABp, confirming differences in promoter specificity towards chemical cues. In addition, more than half of the 28 compounds triggered at least two promoters, confirming vast regulatory cross-talk between the major players of transport in *E. coli* (ED Fig. 2a). Protein synthesis inhibitors emerge as the most promiscuous compounds, as tetracyclines and macrolides triggered simultaneous transcriptional responses in up to 5 out of 7 promoters (Fig 1e).

Looking at individual CPIs, we were able to recapitulate several canonical compound-promoter pairs, including salicylate*-*marRABp^18,24^, paraquat*-*soxSp^17^, procaine-micFp^31^, procaine-ompFp^35^, and 2,2-bipyridyl-micFp^25^, confirming that our screen correctly captures existing knowledge (Fig. 1e and examples at ED Fig. 2b). Compound-dependent repression occurred mostly for the *rob* promoter, consistent with previous reports that *rob* is subject to repression by e.g. MarA and SoxS^36–38^ (Fig. 1c & e). Furthermore, our network shows a rather specific oxidative-stress response of soxSp, also consistent with previous reports^37^. Importantly, our approach misses known interactions, for instance vanillin-marRABp or chloramphenicol-marRABp^13,39^. We attribute this to the facts that we used stringent statistical cutoffs to minimize false positive discovery, and comparatively low concentrations aiming at identifying stronger and specific interactions. Thus, we are most likely underestimating CPIs. Nevertheless, ∼80% of all CPIs we describe here have not been previously reported (Fig. 1d). For instance, we identified tetracycline derivatives, e.g. doxycycline and minocycline, as novel marRABp inducers, beyond the previously reported tetracycline itself^39^. Importantly, we identified macrolides (azithromycin, clarithromycin and erythromycin) and antifolates (sulfonamides and trimethoprim) as novel antibiotic classes triggering several of the tested promoters (Fig. 1e). Beyond antibiotics, we identified the widely consumed food ingredient caffeine as novel marRABp and micFp inducer. Among new inducers, we independently validated that *marRAB* native expression is indeed induced by clarithromycin treatment using RTq-PCR (ED Fig. 2c). In addition, RTq-PCR analysis revealed that also sulfamethoxazole triggers marRABp, although our screening approach was unable to capture it due to stringent cutoffs (ED Fig. 2c).

### General principles driving transport compound-promoter interactions

Capitalizing on our efforts of assessing promoter activity across environmental cues, we next aimed at uncovering general features driving environment-dependent transport regulation in *E. coli*. Since many of our CPIs involved antibiotics, we questioned whether antimicrobial activity is a pre-requisite for modulating gene expression. Indeed, we observed that our set of 28 compounds instigate, on average, lower minimum growth (OD_AUC_) when compared to the compounds that did not trigger the tested promoters (Fig. 2a). However, also compounds without antimicrobial activity at the concentrations tested (e.g. vanillin, caffeine, growth identical to control ay all concentrations tested) are represented among our CPIs, and thus antimicrobial activity is not required to trigger transcriptional changes in transport related genes (Fig. 2a). Next, we asked whether the nature of the antibiotic – bactericidal or bacteriostatic – correlates with its ability to trigger transcriptional changes amid the tested promoters. Interestingly, we found that bacteriostatic antibiotics are enriched for extreme Z-scores (Fig. 2b). In fact, bacteriostatic antibiotics such as tetracyclines, macrolides or sulfonamides account for half of the 53 CPIs described here. Next, because our previous work showed that antagonism is often associated with decreased intracellular antibiotic concentrations^13^, we hypothesized that compounds impacting transport regulation could decrease the concentration of certain antibiotics, and thus exhibit antibiotic antagonism. We combined our results with the data from our previous study, and found that the compounds that are represented in our set of 53 CPIs have indeed a higher chance than other tested compounds to be involved in antagonistic interactions (Fig. 2c).

**Figure 2:**
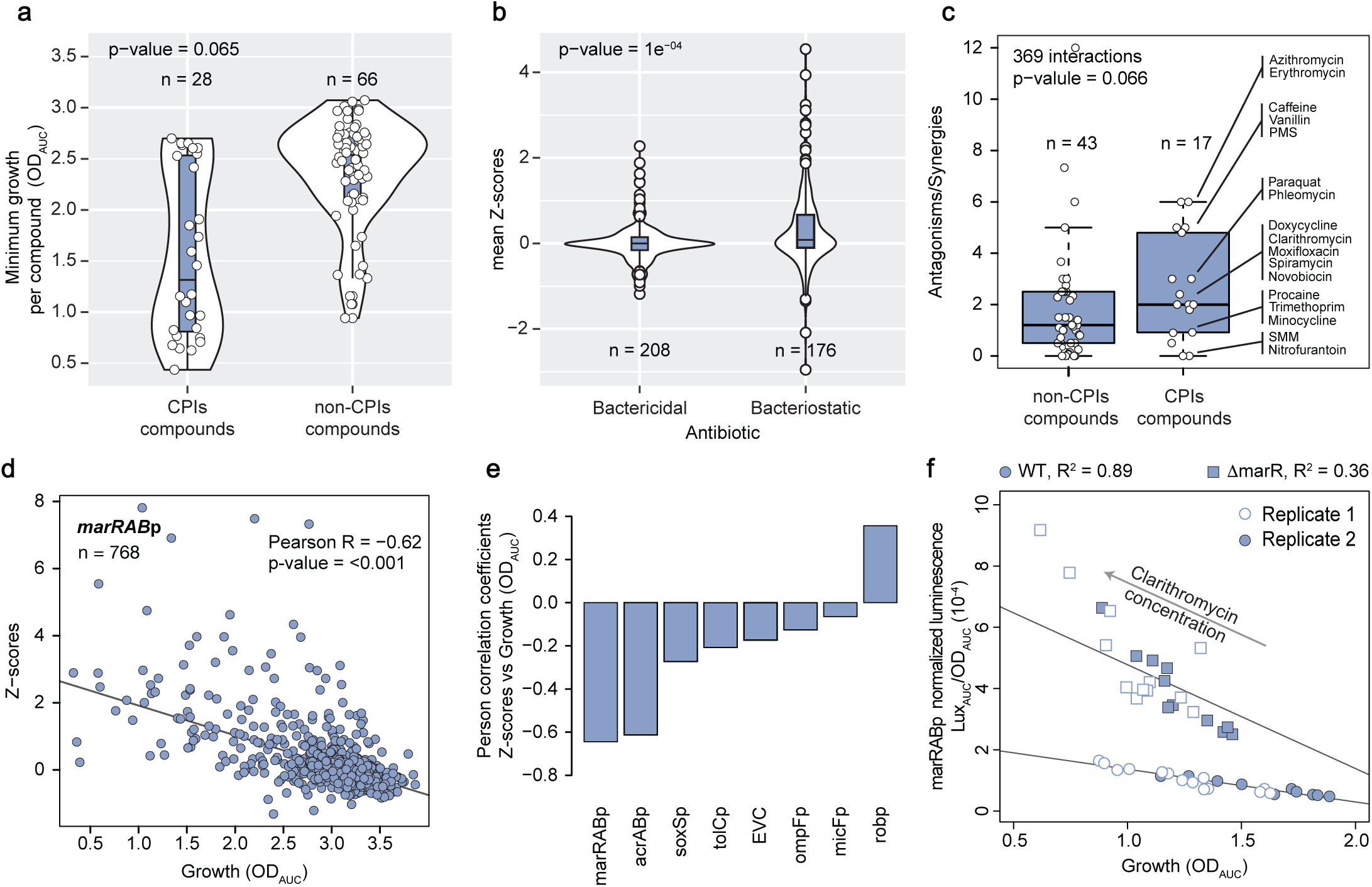
General principles driving transport compound-promoter interactions. **a)** Most compounds identified within CPIs have antimicrobial effect. Distribution of minimum growth (0.1 quantile of all OD_AUC_ measurements for a given compound, 4 concentrations x 8 strains x 2 replicates = 64 values) of all compounds tested (n_total_=94), classified according to whether they are (or not) involved in CPIs. p-value from a double-sided rank sum statistical test shown. Boxplots indicate 25^th^, 50^th^ and 75^th^ percentiles, and whiskers extend up to 1.5 times the interquartile range (IQR) from the 25^th^ and 75^th^ percentiles. **b)** Bacteriostatic antibiotics are over-represented within strong CPIs. Mean Z-scores distributions of all tested compound-promoter pairs involving antibiotics (n_total_=384), classified according to whether the antibiotic is bactericidal or bacteriostatic. p-value from a double-sided rank sum statistical test shown. Boxplots indicate 25^th^, 50^th^ and 75^th^ percentiles, and whiskers extend up to 1.5 x IQR from the 25^th^ and 75^th^ percentiles. **c)** Compounds within CPIs are over-represented among antagonistic drug interactions. Boxplots of ratios antagonisms over synergies (total 369 interactions) for all tested compounds which overlap with our previous work^13^ (n_total_=60), classified according to whether they are (or not) involved in CPIs. p-value from a one-sided rank sum statistical test shown. Boxplots indicate 25^th^, 50^th^ and 75^th^ percentiles, and whiskers extend up to 1.5 times the interquartile range (IQR) from the 25^th^ and 75^th^ percentiles. **d)** marRABp activity inversely correlates with growth. Z-scores of all compound-marRABp tested pairs including water across 4 concentrations and 2 biological replicates (n) are plotted against growth (OD_AUC_). A strong negative linear relationship is illustrated by the line of best fit (Huber robust model). Correlation p-value (double sided t-test) shown. **e)** Promoter activity is generally not correlated with growth. Pearson correlation coefficients of Z-scores versus growth (OD_AUC_) for each individual promoter. A strong negative Pearson correlation is only observed for marRABp and acrABp, while robp shows the opposite behavior. Correlation p- value (double sided t-test) < 0.005 for all promoters. **f)** Induction of marRABp by clarithromycin, as well as its negative correlation with growth, are independent of MarR. Luminescence profiles over growth were measured across a linear range of clarithromycin concentrations from 0 µg/ml to 119.6 µg/ml in wild-type and Δ*marR* background. Growth-normalized luminescence is plotted against growth for two independent biological replicates, and lines-of- best-fit are shown to highlight strong correlation between the two variables.

Driven by the fact that growth inhibition is a strong trait of the compounds represented in the 53 CPIs (Fig. 2a), we probed whether growth alone could explain the extent of promoter induction for any given promoter. By assessing correlation of Z-scores versus growth (OD_AUC_) across all compounds for each individual promoter, we observed strong and significant negative correlation for marRABp and acrABp – meaning that these promoters get progressively stronger activation with increasing inhibitory capacity of the compound at hand (Fig. 2d & e, ED Fig. 2d). This striking observation points towards a generalizable, non-specific, growth-driven response of marRABp across environments, in addition to the widely reported stress-specific response based on de-repression of promoter activity through compound-repressor binding (salicylate-MarR)^26,40^. This finding provides a basis for promiscuous marRABp transcriptional activation by structurally diverse compounds, such as salicylate, clarithromycin and sulfamethoxazole (ED Fig. 2e). We then confirmed that concentration-dependent marRABp transcriptional activation by clarithromycin and sulfamethoxazole indeed remain irrespective of whether its repressor MarR is present or not (Fig. 2f, ED. Fig. 2f). The inverse correlation was observed for robp, where reporter expression increases with growth (ED Fig. 2d). This finding increases the scope of previous observations that MarA may directly or indirectly cause *rob* down-regulation^36,37^, as we find they follow this opposite trend across several environmental conditions. For the remaining promoters no comparable correlation was observed, suggesting growth-independent regulation.

#### Mapping regulator contributions to compound-promoter interactions

To gain more specific insight into the mar-sox-rob network operational modes – in particular into whether and how regulator-promoter hierarchy changes across environmental cues – we generated individual knockout strains of the regulators *marA*, *soxS* and *rob*, and again probed the 658 compound-promoter pairs with our initial chemical library. Promoter preference towards a given regulator (as seen by loss of reporter activity upon regulator deletion) could already be observed for marRABp, soxSp, acrABp and micFp through altered basal promoter activity without any stress (Fig. 3a). In most cases there seems to be a preference for a single regulator, for instance, acrABp activity decreases only when MarA is absent. Nonetheless, micFp seems to be differently controlled, as its activity decreases in the absence of either MarA or Rob, pointing towards cooperation between these two regulators to sustain *micF* basal expression. Our findings add on previous observations of how the different regulators influence promoter activity^37,41^. For instance, MarA not only increases acrABp and micFp activity upon overexpression as it has been shown before^42,43^, but also determines their basal expression in the absence of any chemical or genetic stress (Fig. 3a). In addition, we observed that deletion of either *marA*, *soxS* or *rob* sensitizes *E. coli* towards multiple growth inhibiting compounds, particularly at sub-MIC concentrations (ED Fig. 3a), This highlights that, despite their different responses towards various compounds, the three regulators uniquely contribute to ensure optimal growth.

**Figure 3:**
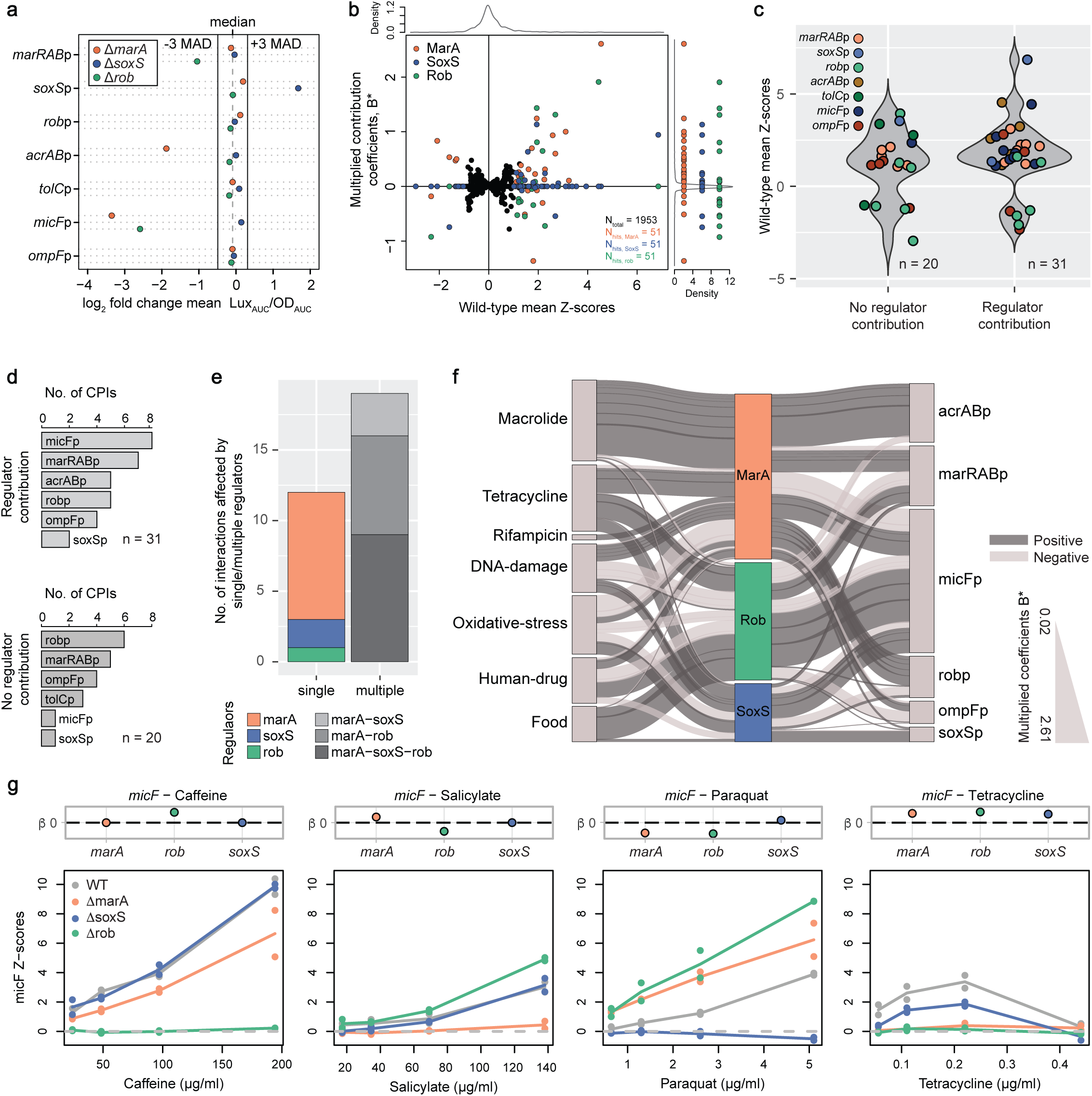
Mapping regulator contributions to compound-promoter interactions. **a)** Deletion of *marA*, *soxS* and *rob* alter promoter basal activity. log_2_ fold-change of water- promoter mean normalized luminescence (Lux_AUC_/OD_AUC_) of each deletion background in relation to the wild-type are plotted. The dashed line represents the median of log_2_ fold-change of water-promoter mean Lux_AUC_/OD_AUC_ across all promoters, and deletion backgrounds. Full lines shown +/3 MAD (). Water-promoter mean Lux_AUC_/OD_AUC_ over 16 replicates per reporter (n = 8 x 2 biological replicates = 16). **b)** Regulator contributions to CPIs are complex and multi- directional. Multiplied coefficients (B*) of MarA, SoxS and Rob of 651 compound-promoter pairs versus wild type mean Z-scores are plotted (n_total_). B* corresponding to 51 significant CPIs are colored according to regulator. Regulator B* are also projected into a single axis (right side) to facilitate visualization. **c)** Most CPIs feature contributions of at least one regulator. Mean Z-score distributions of 51 CPIs (colored by promoter) classified on whether (or not) they have at least one non-zero B*. **d)** Almost all acrABp and micF CPIs depend on MarA, SoxS or Rob. Number of CPIs with (B*≠0, upper plot) and without (B*=0, bottom plot) regulator contributions distributed by promoter. **e)** Most CPIs depend on two or all three regulators. Number of CPIs depending on single of multiple regulators, colored according to which regulators have B*≠0. **f)** Regulator-CPI network. Regulator contributions to 31 CPIs are shown as edges in a Sankey diagram connecting the compounds (left nodes, grouped according to class or purpose) the promoters (right nodes) via the regulators (middle). Edge thickness and node size represent B*, and the total number of interactions, respectively. **g)** Regulator contributions to specific promoters are compound dependent. B* (top) and Z-scores (bottom) of micFp interactions with caffeine, salicylate, paraquat and tetracycline in wild-type and Δ*marA*, Δ*soxS* and Δ*rob*. Lines are colored by strain and indicate mean Z-scores of two biological replicates (dots). Depending on the compound, micFp activity mostly depends on a single (caffeine), on two (salicylate), or on all three regulators (tetracycline and paraquat).

Next, we built a simple machine learning model (Lasso regression for hierarchical interactions^44^) to estimate the contribution of MarA, SoxS and/or Rob to CPIs in our dataset. Briefly, we modeled the transcriptional effect of each compound on a given promoter (interaction scores) as a linear function of compound concentration and regulator presence/absence. The the individual regulator and compound concentration contributions to the observed changes in promoter activity are captured in the model coefficients β_j_ with *j* ∈ {*conc, rob, marA, soxS*} (Methods, ED Fig. 3b). Lasso penalization^45^ was used to obtain sparse model coefficients, and as a result, the majority of the values for each β_*j*_ across all modelled CPIs is 0 (or very nearly 0), with positive and negative deviations reflecting positive or negative contributions to promoter activity in the presence of a given compound (ED Fig. 3c). Firstly, the model accurately captures strong compound concentration dependent effects (reflected by large absolute values for β_#$%#_, ED Fig. 3c), stressing the added-value of concentration-resolved experiments. We also estimated coefficients for synergistic regulator contributions 8: 8_marA,soxS_, 8_marA,rob_ and 8_soxS,rob_. However, we observed that these are much more modest than single regulator coefficients (ED Fig. 3c), and therefore decided to focus on single regulator contributions (β_marA_, β_soxS_, β_rob_) to the 51 CPIs identified in our initial wild-type dataset (2 CPIs, Phleomycin-marRABp and Phleomycin-acrABp, could not be assessed in the regulator deletion mutants due to very poor growth). In order to facilitate interpretation, we computed *multiplied model coefficients* (B*) by multiplying β by the absolute Z-score of the corresponding CPIs in the wild-type. Thus, B* reflect the overall relevance of a regulator towards change in promoter activity by a given compound in the wild-type, since it accounts for the amplitude of the change. B* differ from β in that all B* for weak (or null) compound-promoter interactions scores in the wild type are close to zero (Fig. 3b and ED Fig. 3d). Notably, all regulators were found to have both positive and negative contributions to promoter activity depending on the compound-promoter pair (Fig. 3b), suggesting a high network cross-talk and plasticity towards the environment. Consistent with its rather specific role in response to oxidative stress, SoxS has the tightest zero-centered coefficient B* distribution, with the least number of strong negative or positive contributions. For instance, we accurately capture its positive contribution to auto-regulation during paraquat treatment (paraquat-soxSp, Fig. 3b). MarA emerges as the most prominent regulator across all stresses, with the highest number of non-zero contributions, which are positive in most instances (Fig. 3b). Interestingly, the far less well characterized Rob plays a more prominent role than SoxS across a variety of stresses, with both positive and negative strong contributions to ∼1/3 of all CPIs (Fig. 3b).

Overall, our approach captures contributions of MarA, SoxS or Rob to only 31 out of 51 compound-regulator pairs (Fig. 3c and d). This result is expected, since contributions of other regulatory elements are certainly in place and not taken into account here – e.g. regulation by other transcription factors, such as OmpR or AcrR^2^. Concomitantly, several CPIs involving ompFp and robp – the first does not even contain a mar-sox-rob box – are not influenced by any of the three tested regulators (Fig. 3d). Yet, we quantified MarA, SoxS or Rob contributions to as many interactions involving ompFp, presumably emerging from indirect regulatory network effects. Importantly, the three regulators play a role in pretty much all CPIs involving acrABp and micFp. Among the 31 CPIs for which we could map regulator contributions, only 12 mapped to a single regulator - MarA, SoxS or Rob (Fig. 3e). For the remaining 19 CPIs, they can only be fully achieved when two or all three regulators are in place (Fig. 3e). A more explicit/functional inspection of regulator contributions to CPIs revealed striking observations.

First, the transport-controlling regulatory response to tetracyclines and macrolides is vastly different, despite their common mechanism of action – inhibition of protein synthesis (Fig. 3f). While response to macrolides seems to be strictly driven by MarA, tetracyclines induce a much more complex response involving all three regulators. Another interesting observation is that acrABp expression is almost exclusively controlled by MarA across all compounds tested, with Rob playing a very minor and even negative role (Fig. 3f and ED Fig. 3e & f). Even though MarA control of acrABp is well supported by several studies^42,46,47^, our results indicate that this is a prevalent regulatory relationship across environments, where SoxS and Rob have minor and rather specific contributions. Interestingly, not all compounds triggering marRABp necessary trigger acrABp though, so other factors, such as alternative regulation by AcrR might be determinant in these cases. A very different pattern is observed for micFp, where CPIs are the net outcome of a complex regulatory pattern of positive and negative contributions of MarA, SoxS and Rob (Fig. 3f). We highlight a few examples where micFp expression is strongly influenced by one (caffeine), two (salicylate) or all three (tetracycline and paraquat) regulators (Fig. 3f). Our results so far show how *E. coli* diversifies its response to different chemical stresses and, most importantly, provide a quantitative overview on regulator contribution to promoter activity.

#### Rob-dependent cafieine-micFp interaction causes species-specific antibiotic antagonism in *E. coli*

To showcase the potential of our dataset for uncovering new molecular mechanisms, we chose to validate and further investigate the physiological consequences of caffeine-induced increase of MicF promoter activity – caffeine-micFp interaction (Fig. 1e). Our choice was anchored to two points: first, it involves caffeine - a widely used food ingredient not previously known to impact transport regulation in prominent enterobacteria. Second, because caffeine- micFp interaction emerges from our dataset as the top CPI being primarily/exclusively controlled by Rob (Fig. 3g and 4a), which is by far the least functionally characterized regulator, in comparison to MarA and SoxS. We started out by validating our initial observation that caffeine triggers *micF* expression. Indeed, MicF small RNA levels increase ∼6-fold in the presence of caffeine when compared to no-caffeine control, as measured by Northern blot (Fig. 4b). Next, we confirmed that the levels of OmpF, a major entry point for antibiotics in *E. coli* and a target of MicF, are significantly decreased in the presence of caffeine (Fig. 4c). This result suggests that caffeine could decrease compound uptake (as we previously showed^13^) via upregulation of *micF*, and consequent inhibition of OmpF translation (Fig. 4d). In order to test this hypothesis, we first re-evaluated the outcome of caffeine combination with ciprofloxacin, a fluoroquinolone predominantly taken up through OmpF in *E. coli*, which our previous work showed to be an antagonism^13^. Indeed, a checkerboard assay confirmed the antagonism, showing that the concentration of ciprofloxacin needed to inflict a given inhibitory effect progressively increases with increasing caffeine concentrations (isoboles moving rightward, Fig. 4e). Caffeine alone has no inhibitory effect within the concentrations tested (ED Fig 4). Next, we confirmed that deletion of *micF* or *ompF* completely abolished the antagonism (straight vertical isoboles), as expected, provided that our hypothesis is correct (Fig. 4e). Furthermore, we also observed that this antagonism is strictly Rob-dependent, as *rob* deletion equally abolished the antagonism (Fig. 4e). Altogether, these results fully sustain our hypothesis that caffeine antagonizes ciprofloxacin in *E. coli* via Rob-dependent up-regulation of *micF*, and consequent decrease of OmpF levels Fig. 4d). Importantly, deletion of *marA*, also upregulated by caffeine in a Rob-dependent manner, did not change the caffeine-ciprofloxacin antagonism (Fig. 4f-h), further establishing Rob’s major role in regulating caffeine response in *E. coli*. Interestingly, marRABp upregulation by caffeine did not lead to significant increase in acrABp promoter activity (Fig. 4i), thereby potentially hindering an even stronger antagonism to ciprofloxacin via decreased uptake combined with increased efflux. In summary, our results suggest that caffeine is a novel and specific inducer of Rob transcriptional activity, and this fully explains the molecular mechanism of caffeine-ciprofloxacin antagonism in *E. coli*.

**Figure 4:**
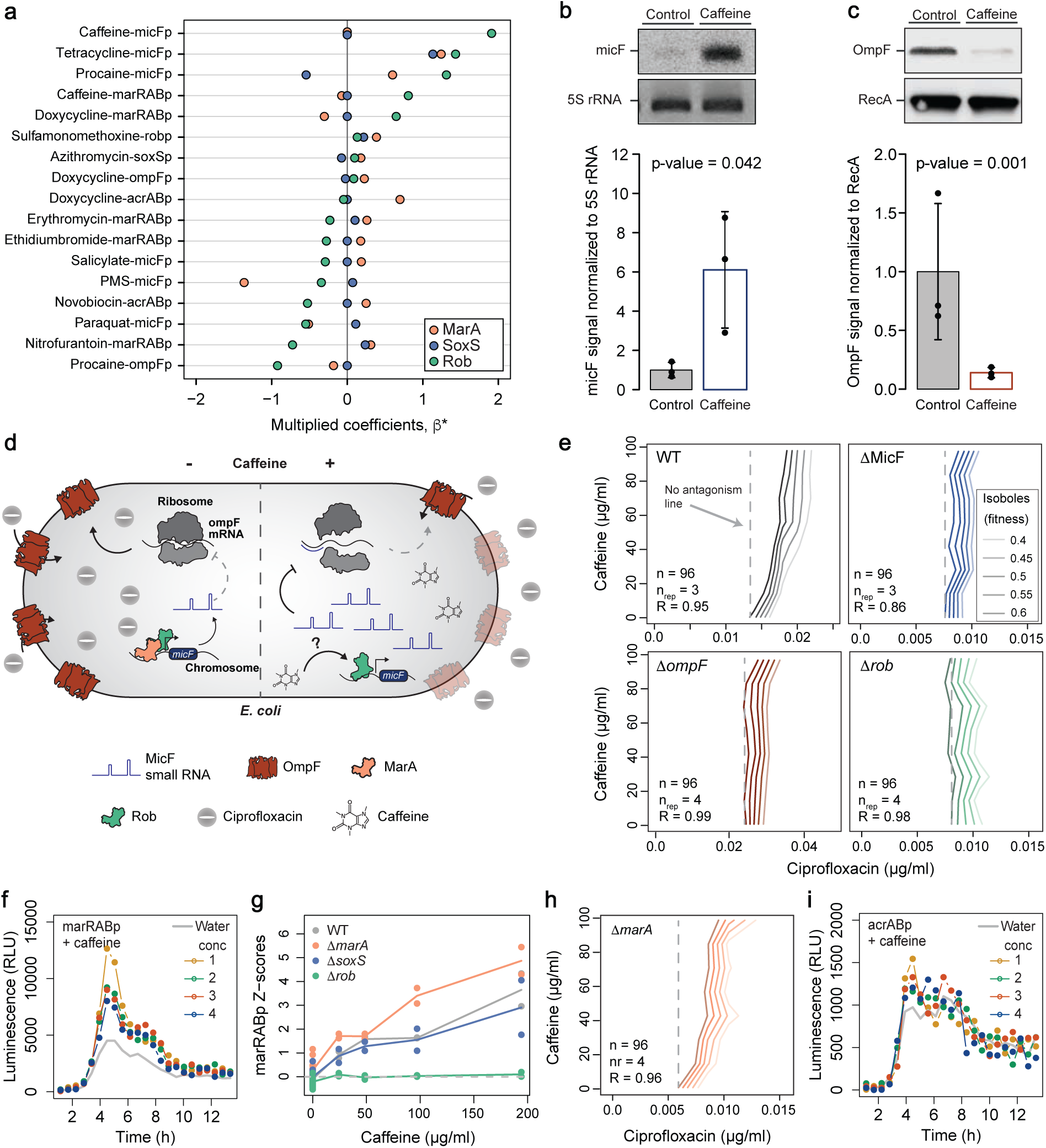
Caffeine induces micFp via Rob causing antagonistic interaction with ciprofloxacin. **a)** Caffeine-micFp interaction is primarily controlled by Rob. β* for all CPIs with non-zero Rob coefficients (n = 17), colored by regulator. **b)** Caffeine increases micF small RNA levels. Northern blot analysis (top) confirming increased levels of small RNA micF upon caffeine treatment (1 mM). One out of three biological replicates (quantification at the bottom) is shown. **b)** Caffeine decreases OmpF protein levels. Immunoblot analysis using whole cell lysate and an *E. coli* OmpF specific antibody shows OmpF decreased levels upon caffeine treatment (1 mM). One out of three biological replicates (quantification at the bottom) is shown. **d)** Proposed model for the molecular mechanism of caffeine-ciprofloxacin antagonism. Caffeine triggers expression of *micF* small RNA in a Rob-dependent manner, which then binds to the 5’-UTR of OmpF mRNA to inhibit and decrease OmpF protein levels. This ultimately prevents ciprofloxacin from entering the cell, resulting in of caffeine-ciprofloxacin antagonism. **e)** Caffeine-ciprofloxacin antagonism in *E. coli* is *micF-*, *OmpF*- and *rob*-dependent. Isobologram for caffeine-ciprofloxacin for *E. coli* wild-type, Δ*MicF*, Δ*ompF* or Δ*rob*. Rightward oriented isoboles indicate antagonism, while upward oriented isoboles indicate no antagonism. A dashed line is plotted for no-antagonism reference for isobole 0.6 for all strains. One out of 3 or 4 biological replicates (n_rep_) is shown. R is the Pearson correlation between the biological replicates obtained with 96 (n) fitness values used to obtain each checkerboard. **f & g)** Caffeine induces marRABp in a Rob-dependent manner. f) Luminescence (RLU) profiles of marRABp basal activity (grey) and with increasing concentrations of caffeine (conc, Supplementary table 2) over time are shown. One out of two biological replicates is shown. g) Z-scores of caffeine-marRABp interaction showing its dependency on rob. Lines are colored by strain and indicate mean Z-scores of two biological replicates (dots). **h)** Deletion of marA does not affect the antagonism between ciprofloxacin and caffeine. Isobologram for caffeine- ciprofloxacin for *E. coli* Δ*marA*. Details as in panel e. **i)** Caffeine causes marginal or no- induction of acrABp. Luminescence (RLU) profiles of acrABp basal activity (grey) and with increasing concentrations of caffeine (Supplementary table 2) over time are shown. One out of 2 biological replicates is shown.

Interestingly, our previous work showed that antagonisms involving caffeine are invariably specific to *E. coli*, as they were not found for the closely related species *Salmonella enterica* Typhimurium^13^. Indeed, also here we could confirm that caffeine does not change ciprofloxacin activity in *S.* Typhimurium (Fig. 5a). Also here, caffeine alone does not inhibit the growth of *S.* Typhimurium (ED Fig. 4). Even though the mar-sox-rob box, as well as all three regulators, are also present, they could operate very differently, as *S.* Typhimurium contains yet an additional regulator, RamA^2^. Nonetheless, OmpF posttranscriptional regulation by MicF is well documented for *S.* Typhimurium. Thus, the reason for lack of conservation of the antagonism between the two species must be upstream of MicF, e.g. different regulation, or downstream of OmpF production, e.g. alternative antibiotic transport. We first confirmed that caffeine also increases micFp expression in *S.* Typhimurium, using a lux-reporter plasmid for micFp from *S.* Typhimurium similar to that used in *E. coli* (Fig. 5b). Furthermore, we observed that caffeine treatment also reduces the level of OmpF in *S.* Typhimurium as quantified by Western blot (Fig. 5c). Altogether, this shows that caffeine-induced regulatory effects are conserved between the two species, and yet the antagonism is only present in *E. coli*. This observation confirms that OmpF-mediated uptake is most probably not determinant for ciprofloxacin activity in *S.* Typhimurium to the same extent it is for *E. coli*, as previously suggested^48^. Indeed, we could confirm that *ompF* deletion alters ciprofloxacin sensitivity in *E. coli*, but not in *S.* Typhimurium (Fig. 5c and d). Several reasons could explain why this is the case, for instance the presence and alternative regulation of additional outer membrane porins in *S.* Typhimurium, or a more prevalent role of efflux versus uptake^48–50^. Nonetheless, these results ultimately illustrate how challenging it is to predict final outcome of CPIs on drug transport across species, even closely related ones.

**Figure 5:**
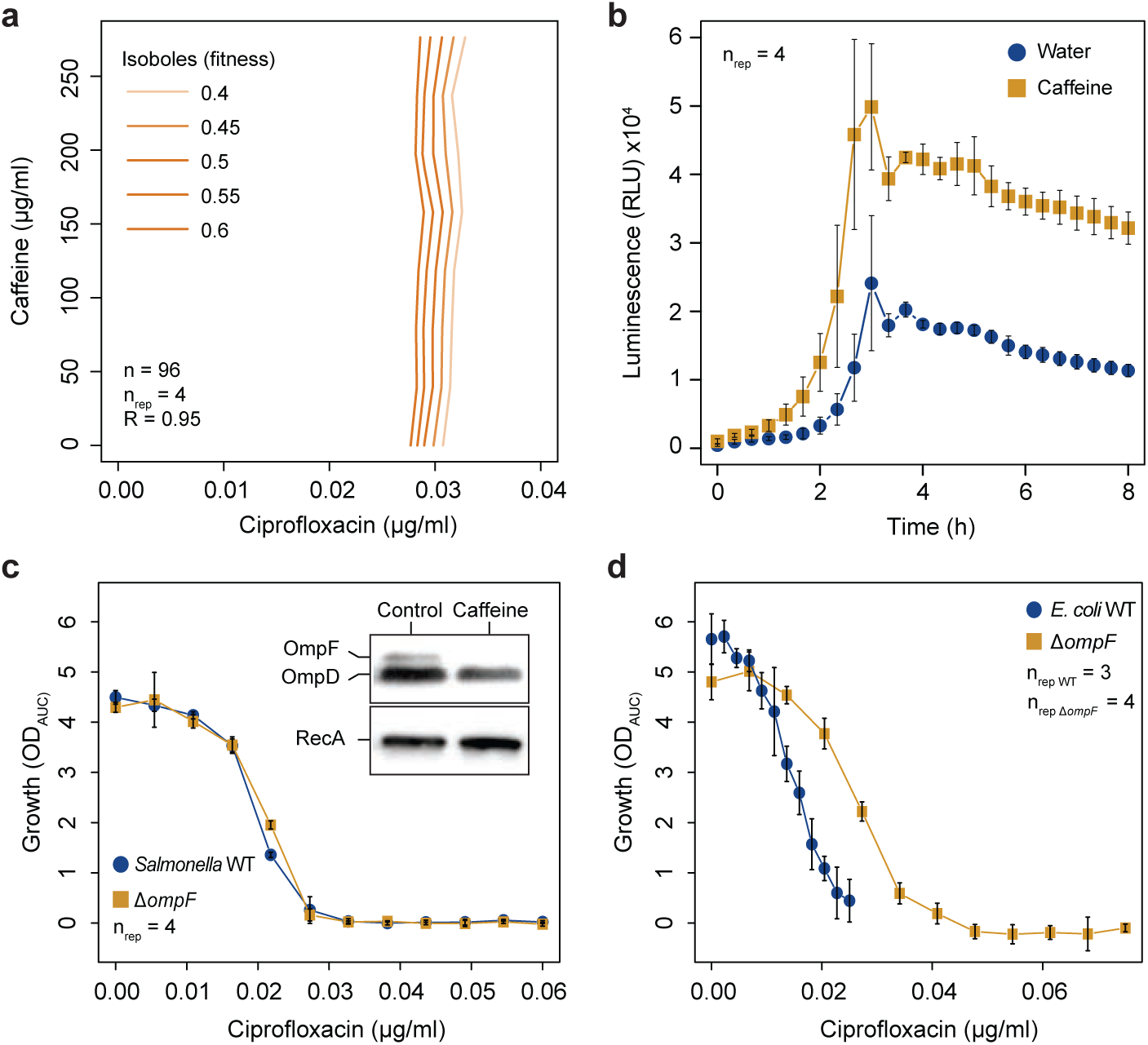
Caffeine-ciprofloxacin antagonism is absent in *S.* Typhimurium despite conserved regulatory mechanism. **a)** Absence of caffeine-ciprofloxacin in *S.* Typhimurium. Isobologram for caffeine-ciprofloxacin for *S.* Typhimurium. Details as in Fig. 4e. **b)** Caffeine induces micFp activity in *S.* Typhimurium. Luminescence profiles of *S.* Typhimurium micFp reporter strain over time +/- caffeine (2 mM). Mean luminescence (dots) and standard deviations (error bars) over 4 biological replicates. **c)** Ciprofloxacin MIC is not altered upon OmpF deletion in *S.* Typhimurium. Ciprofloxacin MIC curves (growth vs antibiotic concentration) of *S.* Typhimurium wild-type and Δ*ompF.* Mean growth (OD_AUC_, dots) and standard deviations (error bars) over 4 replicates. Inlay: Immunoblot analysis using total protein extractions and an OmpF polyclonal antibody shows OmpF decreased levels upon caffeine treatment (2 mM). One out of 3 biological replicates is shown. **d)** Ciprofloxacin MIC decreases upon OmpF deletion in *S.* Typhimurium. Ciprofloxacin MIC curves (growth vs antibiotic concentration) of *E. coli* wild-type and Δ*ompF.* Mean growth (OD_AUC_, dots) and standard deviations (error bars) over 3 or 4 biological replicates.

## Discussion

Transport across the bacterial cell envelope has been subject of study for decades, with important constituents like AcrAB-TolC and OmpF, and also central regulators such as MarA, SoxS and Rob being extensively characterized^2,15^. Despite vast molecular evidence on how each of these proteins plays a role in a complex regulatory cascade, we mostly lack comprehensive approaches that provide general overviews on how all the different players come together to orchestrate the coordinated response. Here we provide an integrative systematic approach to assess transcriptional response of transport genes to environmental cues (CPIs), while quantitatively assessing regulator contributions to each response. Such an approach enabled not only the discovery of new CPIs, such as caffeine-micFp and macrolides- marRABp, but most importantly, allowed us to disentangle the relative contribution of each central regulator to the observed phenotype. Our results endow many several previously postulated concepts, such as that not all compounds being effectively effluxed^2,3,51^, for instance the bactericidal antibiotics fluoroquinolones and β-lactams are capable of inducing expression of efflux genes, or that transport regulation underlies antibiotic antagonism^13^. Importantly, we provide a set of new general findings suggesting a paradigm change in our perception of how *E. coli* controls its transport. For instance, *marA* regulation is mostly investigated in the presence of salicylate, which releases *marA* transcriptional repression by binding MarR^26^. However, our data shows that this represents a rather specific case, and that most compounds seem to regulate *marA* in a growth dependent but MarR independent manner. Another interesting finding is that ∼1/3 of all CPIs depend on Rob. This is well beyond what is generally described or perceived, and indicates that a much better understanding Rob- dependent regulation is crucial to fully comprehend transport transcriptional control. Interestingly, to almost all CPIs towards which Rob contributes, it does not do so alone, but mostly in tandem with MarA, or MarA and soxS (Fig. 3e). Thus, the reason why its role has been long overlooked is perhaps the fact that most available studies on this matter focus on one of the three regulators alone (MarA, SoxS or Rob). Exceptionally, our data shows that Rob alone seems to have complete control (in comparison to MarA and SoxS) of transcriptional effects of caffeine at comparatively lower concentrations to previously described “rob-inducing compounds”, such as dipyridyl or bile-acids^16,25^. Last, our approach provides very clear evidence and quantification of highly coordinated transcriptional control towards different environments, even though most of the tested promoters contain a mar-sox-rob box: promoter transcriptional activity and regulator contributions to it is highly dependent on the environmental cue, as we could show, for instance, by profoundly different regulation of marRABp, acrAB and micF. Nonetheless, we acknowledge limitations in our study both in the degree of chemical space and promoters screened, and in that we provide only transcriptional output, without information on posttranscriptional regulation or alterations on a translational level. Expanding future endeavors in these directions will unequivocally add on our current findings.

A second aspect to highlight is the potential of our dataset for mining molecular mechanisms. We showcased it by exposing how caffeine-micFp interaction underlies caffeine-antibiotic antagonism through impaired antibiotic uptake, in a Rob-dependent manner. This finding features the importance of the immediate environment for treatment efficacy through transport modulation, and supports future predictive design of drug combinations. Given the ever increasing evidence of transport’s prevalent role in reacting to harsh environments, for instance in modulating the impact of human-targeted-drugs in gut microbes^10^, or potentially contributing to bioaccumulation within the gut thereby modulating drug action on the host^11^, the scope of our findings and approach goes well beyond *E. coli* or even enterobacteria. Nonetheless, our observation on lack of conservation of drug transport mechanisms even between *E. coli* and the closely related *S.* Typhimurium leads us to foresee a challenging, but unavoidable and important task in mapping key determinants of transport functions across different bacteria.

## Acknowledgments

We thank Tobias Bergmiller (University of Exeter, UK) and Libera Lo Presti (Tübingen University, GER) for providing feedback on the manuscript, and the Brochado lab members and CB thesis advisory committee for discussions. We thank Franziska Faber and Manuela Fuchs (University of Würzburg, GER) for advice with Northern blot. We thank Takuya Shiota and Trevor Lithgow (Monash University, AUS) for sharing the *E. coli* specific OmpF antibody. This work was supported by the Deutsche Forschungsgemeinschaft (DFG, German Research Foundation) under Germanβs Excellence Strategy – EXC 2124 – 390838134, and the Emmy Noether program to ARB (GO 3161/1-1). CB and ROA were supported by the StressRegNet consortium awarded to ARB and CM within the Bavarian research network bayresq.net funded through the Bavarian State Ministry of Science and Arts, Germany.

## Author contributions

CB and ARB conceived and designed the study. CB and MW performed the experiments. ROA, MS and CM designed and implemented the regression model. CB and ARB analyzed the data and wrote the manuscript. ROA and CM contributed to manuscript writing. ARB supervised the study.

## Data and Code Availability

The data supporting the findings of this study will be available as source data & supplementary files/Github. All code will be made available on GitHub.

## Methods

### Growth medium, reporter plasmids and strain construction

A summary of all strains used in this study can be found in supplementary table 1. *Escherichia coli* BW25113 and *Salmonella enterica* subsp. *enterica* ser. Typhimurium 14028S were used as wild-type strains (WT) and cultured in Lysogeny Broth (LB Lennox) adjusted to pH 7.5 at 37°C. The medium was supplemented with kanamycin (50 µg/ml, CatNo. K1876-5G, Sigma- Aldrich-Merck), carbenicillin (100 µg/ml, CatNo. Cay20871-5, Biomol) or spectinomycin (100 µg/ml, CatNo. S4014-5G, Sigma-Aldrich-Merck) when selection was required for strain construction.

All plasmids constructed in this study are listed in supplementary table 1. Promoter regions (500 bp upstream of respective start-codon) of interest were amplified from genomic DNA of *E. coli* BW25113 or *Salmonella* Typhimurium using Q5 polymerase according supplier instructions (CatNo. M0491S, New England Biolabs (NEB), USA). All DNA oligos used in this study are listed in supplementary table 1. Reporter plasmids for *E. coli* were assembled by restriction-ligation using enzymes EcoRI and XhoI (SalI in case of acrABp, CatNos. R3101S, R0146S, R3138S, NEB) to linearize the backbone vector pTU175-LUX. T4 DNA ligase (CatNo. M0202S NEB) was used for plasmid assembly following supplier instructions. Reporter plasmids for *S.* Typhimurium were assembled using Gibson Assembly, using XhoI for plasmid linearization. Vector pTU175-LUX is a low copy plasmid with a pSC101 origin of replication constructed from pTU175^1^. by insertion of an oriT, a spectinomycin resistance cassette and the full luxCDABE operon (amplified from Plasmid #44918, AddGene) with a putative RBS for basal expression – used as **e**mpty **v**ector **c**ontrol, EVC. The pASCOT-LUX vector is a variant of pTU175-LUX with a carbenicillin resistance cassette instead of spectinomycin used for *Salmonella*. DNA inserts were digested with the respective Restriction enzymes and assembled into the plasmid backbone utilizing T4 DNA ligase according to a standard protocol (NEB). Plasmids were transduced into *E. coli* BW25113 and *Salmonella* Typhimurium 14028S by transformation (TSS transformation) and electroporation, respectively. All strains and plasmids constructed during this study are available from the authors upon request.

Deletions of Δ*marA*, Δ*soxS* and Δ*ompF* were derived from the KEIO collection in case of *E. coli*^2^ or a similar knockout-library in case of *S.* Typhimurium^3^. Mutations were confirmed using PCR and transduced into wild type BW25113 or ST14028s using P1 and P22 phage, respectively. Deletion of *rob* in *E. coli* was done with Lambda RED recombineering according to the protocol of Datsenko and Wanner^4^, similar to how the KEIO collection strains were created. Kanamycin resistance cassettes were subsequently removed using the pCP20 plasmid.

### Compound-promoter screens

The compound-promoter screen for *E. coli* BW25113 (wild type, as well as Δ*marA*, Δ*soxS* and Δ*rob* backgrounds) containing each reporter plasmid – arcABp, marRABp, soxSp, robp, micFp, ompFp, tolCp and EVC - was done in black 384 well-plates with clear bottom (CatNo. 781097, Greiner-Bio One, Germany), in 40 µl LB Lennox. The compound library contains 94 diverse compounds purchased from Biomol (Germany), MP Biomedicals (Germany), or Sigma-Aldrich-Merck (solvents, concentrations and purchase details listed in supplementary table 2). The highest screening concentration of compounds with antimicrobial activity was adjusted to MIC for antimicrobials, 500 µM for most non-antimicrobials, and up to 1 mM for small compounds with similarity to canonical inducers (positive controls, e.g. salicylate, supplementary table 2). Four working concentrations were achieved through 1:2 serial dilutions using a Biomek i7 liquid handler (Beckman Coulter, ED. Fig. 1a). Precultures were grown overnight and diluted to an OD_600 nm_ of 1 (WT) or 0.1 (deletion backgrounds) and used to inoculate 384 well plates using a Singer Rotor+ replicator (Singer Instruments, UK), resulting in further ∼1:600 dilution and starting OD of ∼0.003 and ∼0.0003. Transparent breathable membranes (Breathe-Easy®, Sigma-Aldrich-Merck) were used to seal plates, which were then incubated at 37°C, shaking at 800 rpm using a Cytomat 2 incubator (Thermo Scientific). Growth (OD_600 nm_) and reporter activity (luminescence) were measured in 30-minute intervals over 12 hours in a Synergy H1 plate reader (Agilent, USA). The screen was performed in biological duplicates, resulting in 768 growth and luminescence curves per strain.

Data analysis was performed using R (version 4.2.2). A representation of the analysis pipeline can be found in ED. Fig. 1b. Baseline-correction of growth curves was done by subtraction of initial OD_600 nm_ before growth onset (time 1-2 hours). Area under the curve was calculated for growth (OD_AUC_) and luminescence (Lux_AUC_) curves between 0 and 8 h in case of the wild type and between 1 and 9 h for the deletion background strains to account for the differences in inoculum size mentioned above. This eight-hour interval covers lag phase, exponential phase and transition into stationary phase assuming regular, non-stressed growth – using water instead of any compound. Non-growing samples (less than 10% OD_AUC_ of water controls) were removed from further analysis (250-419 wells out of 6144 wells total, depending on strain background), and luminescence was then normalized by growth (Lux_AUC_/OD_AUC_, supplementary table 3). Based on the premise that most compounds in the library do not induce/inhibit expression of any of the promoters, compound-promoter interactions were defined to be the deviation (residuals) of the line-of-best fit (Huber robust linear regression)^5^ of normalized luminescence between a given promoter and the EVC. This method allows to better control for possible non-specific transcriptional effects of each compound, as we observed that normalized luminescence for EVC can in fact change across compounds/concentrations (supplementary table 3). Importantly, we observed that the robust linear fits had reasonably high R^2^ (coefficient of determination, supplementary table 4), indicating that this approach captures and corrects well non-specific effects, and positive and negative deviations (residuals) reflect compounds which increase or decrease promoter expression, respectively. The interaction scores (residuals) were subsequently Z-transformed (Z-scores) to allow comparability of promoters of varying signal intensity. Finally, significant compound-promoter interactions were called based a double cut-off on mean Z-score of all compound concentrations and replicates for each compound-promoter pair (±1) and Benjamini-Hochberg^6^ adjusted double sided rank-sum test p-value (<0.05) comparing the Z- scores distributions of each compound-promoter to water-promoter.

### DNA and RNA quantification by qPCR and RT-qPCR

To confirm plasmid copy number stability after treatment with protein biosynthesis inhibitors, overnight cultures of *E. coli* BW25113 harboring plasmids pTU175-Lux-EVC or pBR322 were diluted 1:100 in fresh LB medium and grown at 37°C with agitation until exponential phase (OD _600 nm_ ∼0.4). Chloramphenicol was added to half of the cultures to a final concentration of 2 µg/ml followed by further cultivation for 30 minutes. Total DNA was extracted using the Monarch Genomic DNA Purification Kit (NEB) using the manufacturer’s instructions. All experiments were conducted in three biological replicates. Relative plasmid number fold changes were estimated by comparison with a non-treated control. All DNA oligos used in this study are listed in supplementary table 1.

To confirm induction of *marA* after treatment with salicylate clarithromycin and sulfamethoxazole, overnight cultures of wild-type *E. coli* BW25113 were diluted 1:100 in fresh LB medium and grown at 37°C with agitation until exponential phase (OD _600 nm_ ∼0.4). Salicylate, clarithromycin and sulfamethoxazole were added to cultures to a final concentration of 1 mM, 40 µM and 0.8 mM, respectively, while controls were left untreated, followed by further cultivation for 30 minutes. Total RNA was extracted using the Monarch Total RNA Miniprep Kit (NEB) using the manufacturer’s instructions. Luna Universal One-Step RT-qPCR Kit (NEB) was used to prepare cDNA and as reagent for RT-qPCR according to the manufacturer’s instructions. All experiments were conducted in at least three biological replicates and relative expression levels were estimated as previously described^7^, using *gyrA* expression as reference.

### Lasso regression for estimation of regulator contributions of compound-promoter interactions

Firstly, the interaction scores gathered from the different genetic backgrounds were quantile normalized to account for baseline changes in expression due to regulator deletions. These normalized scores were then further pre-processed to consider the baseline values obtained from exposure to water through a soft-thresholding approach. Namely, we subtract the value of the water-promoter score from compound-promoter scores gathered in the same genetic background. In cases where the compound-promoter score is lower than the corresponding water-promoter score, the compound-promoter score is set to 0. These normalized and *water- thresholded* scores are then centered and scaled prior to modelling.

Once pre-processed, we modelled the scores for a given CPI as a function of compound concentration and genetic background, resulting in a design matrix *X* = [*X_conc_, X_rob_, X_marA_, X_soxS_*] with dimensions *n* x 4 (*n* being the number of samples). Compound concentration is a discrete variable (*X_conc_* ∈ {1,2,4,8}^*n*^), while genetic background is represented as a binary variable to indicate regulator presence/absence (*X_rob, X_marA_, X_sosX__* ∈ {0,1}^*n*^). We also include all pairwise interactions between the variables in *X* in our model. In this way, our model can be stated as:

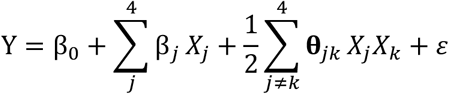

where *Y* ∈ ℝ^*n*^ is the vector of pre-processed interaction scores for a given CPI, β_0_ is a CPI- specific intercept, β_*j*_ is the effect that variable *j* ∈ {*conc, rob, marA, soxS*} has on *Y*, θ_*jk*_ captures the pairwise interaction effect between variables *j* and *k*, where *j*, *k* ∈ {*conc, rob, marA, soxS*} and *j* ≠ *k*, and χ models technical and biological noise.

To select for a parsimonious model, we estimate model coefficients via regularized maximum- likelihood estimation with lasso penalization^8,9^. Furthermore, we restrict non-zero interaction terms θ_*jk*_ to only be present if both associated individual effects β_*j*_ and β_*k*_ are also non-zero (strong hierarchy). This restriction prioritizes explaining *Y* in terms of main effects β, and interactions are only included if the response cannot be solely captured by linear additive effects.

Given that the log-likelihood of our model is 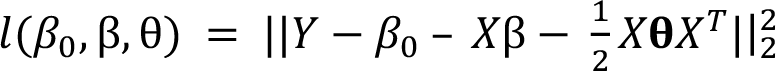, the complete optimization problem for the hierarchical interaction model is:

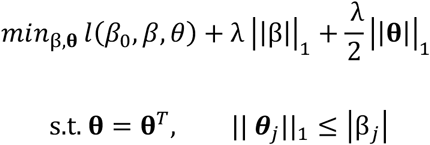

where λ > 0 is the lasso penalization parameter that controls the sparsity of coefficients β and θ.

We solve this optimization problem using the efficient implementation provided in the R package hierNet (version 1.9)^9^. The optimal value for λ was determined through 4-fold cross validation. We selected the λ value that was within one standard error of the λ that minimized the cross validation error. In this way, the nature of the individual effects that drug concentration and regulator presence have on changes in gene expression in response to a given chemical stressor are captured in the sign and magnitude of the coefficients β.

Interaction coefficients θ, potentially reflecting added synergistic effects, were negligible compared to concentration or single regulator coefficients, and were therefore excluded from further analysis.

Regulator contribution coefficients to compound-promoter pairs were then multiplied by the absolute of mean Z-scores of the wild type – multiplied coefficients B* – thereby reflecting the strength of the compound-promoter interactions in wild type, in addition to regulator contribution. This approach facilitates interpretation and enables better understanding of regulator contributions to strong compound-promoter interactions in the wild type.

### Determination of Minimum Inhibitory Concentration (MIC)

Compound Minimum Inhibitory Concentration (MIC) against *E. coli* and *Salmonella* strains was quantified using liquid growth assays in LB-Lennox in microtiter plates in preparation for screening or checker board assays. The assay was performed in 96-well plates in 100 µl, and conditions similar to those used in the screening approach described above. Drugs were diluted linearly in 11 equal steps, allowing for finely resolved quantification of antimicrobial efficacy. Growth curves were analysed similar to those from the screening approach, and growth after 8h was approximated by OD_AUC_.

### Quantification of OmpF levels by Western Blot

*E. coli* and *S. enterica* OmpF levels were quantified by Western blotting. Caffeine treated and untreated cultures were prepared in three biological replicates each by 1:100 dilution of over- night cultures in 20 mL LB-medium and grown at 37°C and 180 rpm shaking. Caffeine was added to half of the flasks at a concentration of 1 mM, while the other half of the flasks served as a negative control. Cultures were grown until OD_600_ of ∼0.8, followed by pelleting via centrifugation (10 min, 4000g, 4°C). Pellets were resuspendet in 200 µl 4% SDS solution, heated to 95°C for 5 min and stored at -80°C until used. The Pierce™ BCA Protein Assay Kit (Fisher Scientific, Germany) was used to calculate total protein concentration of all samples according to the manufacturer’s instructions. Samples were adjusted to 4 µg/µl protein with 4% SDS solution and mixed in equal parts with 2x Laemmli buffer containing 2- mercaptoethanol. Aliquots containing 20 µg of total protein (10 µl) were boiled for 5 min at 95°C to denature any proteins. Samples were separated by SDS-PAGE on a 10% gel at 80 V for 3.5 h and transferred to a PVDF membrane via Trans-Blot® Turbo™ system (BioRad), according to the manufacturer’s instructions. A specific rabbit αOmpF antibody (kindly provided by Joel Selkrig, Aachen, Germany) and a rabbit αRecA antibody (ab63797, Abcam) were used as primary antibodies at dilutions of 1:20000 and 1:5000, respectively, to detect OmpF and RecA on the PVDF membrane. For both primary antibodies, we used an HRP- coupled secondary antibody (A0545, Sigma), and Pierce™ ECL Western Blotting-Substrate (Thermo Scientific) to visualise OmpF using an chemiluminescence imager (Intas Science Imaging Instruments GmbH, Germany).

### Quantification of MicF small RNA by Northern blotting

Bacterial cultures were grown to exponential phase (OD_600_ ∼0.4) and treated with 1 mM caffeine for 30 minutes or left untreated as negative control. Samples were then mixed with 0.2 volumes of STOP solution (95% ethanol, 5% phenol) and snap-frozen in liquid nitrogen to prevent RNA degradation. Total RNA was extracted using the hot phenol method. Cell pellets were thawed and resuspended in 65°C lysis buffer (40 mM EDTA pH 8, 200 mM NaCl, 0.5% SDS, 100 mM Tris-HCl pH 7.5), incubated for 5 minutes at 65°C in a water bath and subsequently mixed with acidic phenol (ROTI Aqua-Phenol, Roth). Samples were mixed thoroughly by vortexing, snap-frozen in liquid nitrogen and centrifuged for 10 minutes. The upper aqueous phase was then mixed with the same volume of chloroform-isoamyl alcohol (24:1) and mixed again by vortexing. The resulting upper phase was then mixed with 1/10^th^ volume of 3M sodium acetate (pH 4.5) and one volume isopropanol to precipitate total RNA for 30 minutes on ice. Supernatants were subsequently removed, pellets dried and resuspended in RNAse free water. Northern blotting, radioactive labelling of DNA oligonucleotides, hybridization and signal detection were all performed as previously described^10^ (ref). Signals were subsequently analysed using ImageJ software^11,12^ (ref).

### Checkerboard Assays

Quantification of drug interactions was performed using checkerboard assays. In brief, a checkerboard assay resembles a two-dimensional MIC assay, with two different drugs being combined across concentration gradients. The assays were performed in 384-well plates in biological quadruplicates and conditions similar to the screening described above. Growth inhibitory effect (OD_AUC_ after 8h) was determined in a series of 7 (vertical dilution series) equally spaced concentrations for caffeine and 11 (horizontal dilution series) equally spaced concentrations for ciprofloxacin. Concentrations were adapted for *E. coli*, *S. enterica*, and respective deletion mutants (Δrob, ΔmicF, ΔmarA and ΔompF). Fitness was calculated by normalization of OD_AUC_ of each well with the no-drug control. Lines of equal fitness (isoboles) were estimated by the contours derived from drug-interaction-surfaces. Provided that caffeine alone does not show inhibitory effects at the concentrations tested, antagonism with ciprofloxacin is reflected by increased concentrations of ciprofloxacin needed to inflict a given inhibitory effect with increasing concentrations of caffeine (isoboles moving rightward, Fig. 4e).

---

**Extended data figure 1:**
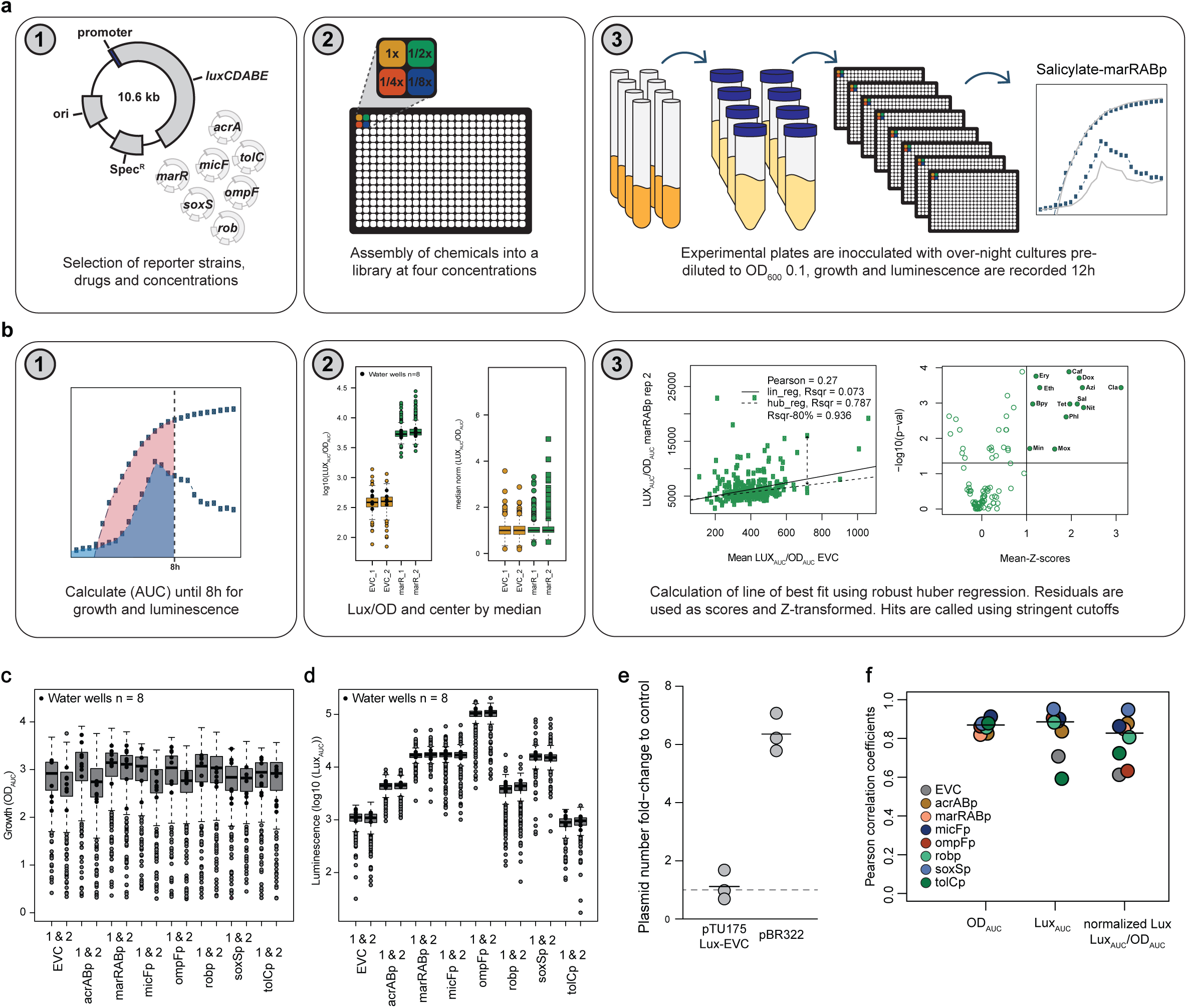
Schematic representation of the screen workflow and data processing. **a)** Schematic screen workflow. Details described in methods. **b)** Schematic of data processing. Details described in methods. **c)** Boxplots of growth (OD_AUC_) across all reporters and replicates. Each boxplot represents a 384 well-plate (n=384). Negative controls (water treatment) are displayed in black (n = 8 per strain). 1 & 2 refer to biological replicates. **d)** Boxplots of luminescence (AUC_LUX_) data across all reporters and replicates. Each boxplot represents a 384 well-plate (n=384). Negative controls (water treatment) are displayed in black (n = 8 per strain). 1 & 2 refer to biological replicates. **e)** Treatment with protein biosynthesis inhibitors does not affect copy number of pTU175 plasmids. Relative fold-change of pTU175- Lux-EVC and pBR322 after treatment with 2 µg/ml chloramphenicol compared to a negative control using qPCR. Three biological replicates are shown. **f)** Pearson replicate correlation of growth (OD_AUC_), luminescence (LUX_AUC_) and normalized luminescence (LUX_AUC_/OD_AUC_) between the duplicates of each strain.

**Extended data figure 2:**
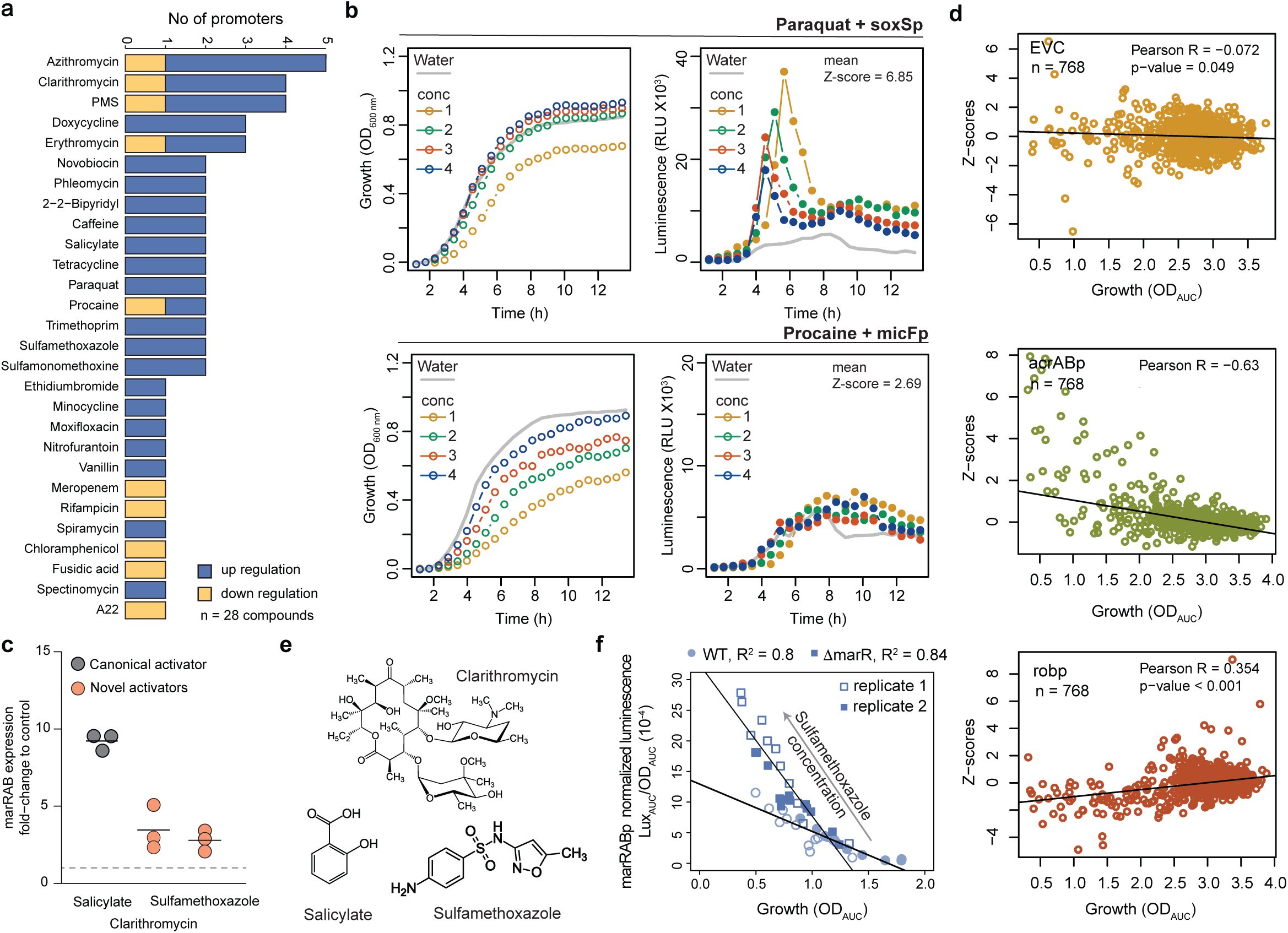
General principles driving CPIs in *E. coli*: additional supporting findings. **a)** General features of CPIs: number of CPIs per compound classified as up and down- regulation. **b)** Previously known CPIs paraquat-soxSp^17^ and procaine-micFp^31^ are captured by our screening approach. Growth (OD_600 nm_) and luminescence (RLU) profiles over time for soxSp (top) and micFp (bottom) basal activity (grey) and with increasing concentrations of paraquat or procaine, respectively (conc, Supplementary table 2) are shown. One out of two biological replicates is shown. **c)** Clarithromycin and sulfamethoxazole are novel inducers of marRABp expression. RNA levels of *marA* after treatment with salicylate (positive control), clarithromycin and sulfamethoxazole. Data was double normalized to a non-treated control and to the house-keeping gene *recA* (Methods). Three biological replicates are shown. **d)** Correlation between growth and promoter activity for EVC, acrABp and robp. Z-scores of all compound-EVC/acrABp/robp tested pairs including water across all 4 concentrations and 2 biological replicates (n) are plotted against growth (OD_AUC_). Pearson correlation coefficients (R) indicate no-, negative and positive correlation for EVC, acrABp and robp, respectively. Correlation p-value (double sided t-test) shown. Linear relationships are illustrated by lines of best fit (Huber robust model). **e)** Chemical structures of known and novel marRABp inducing compounds. **f)** Induction of marRABp by sulfamethoxazole, as well as its negative correlation with growth, are independent of MarR. Luminescence profiles over growth were measured across a linear range of sulfamethoxazole concentrations from 0 µg/ml to 101.2 µg/ml in wild-type and Δ*marR*. Growth-normalized luminescence is plotted against growth for two independent biological replicates, and lines-of-best-fit (pooled replicates) are shown to highlight strong correlation between the two variables.

**Extended data figure 3:**
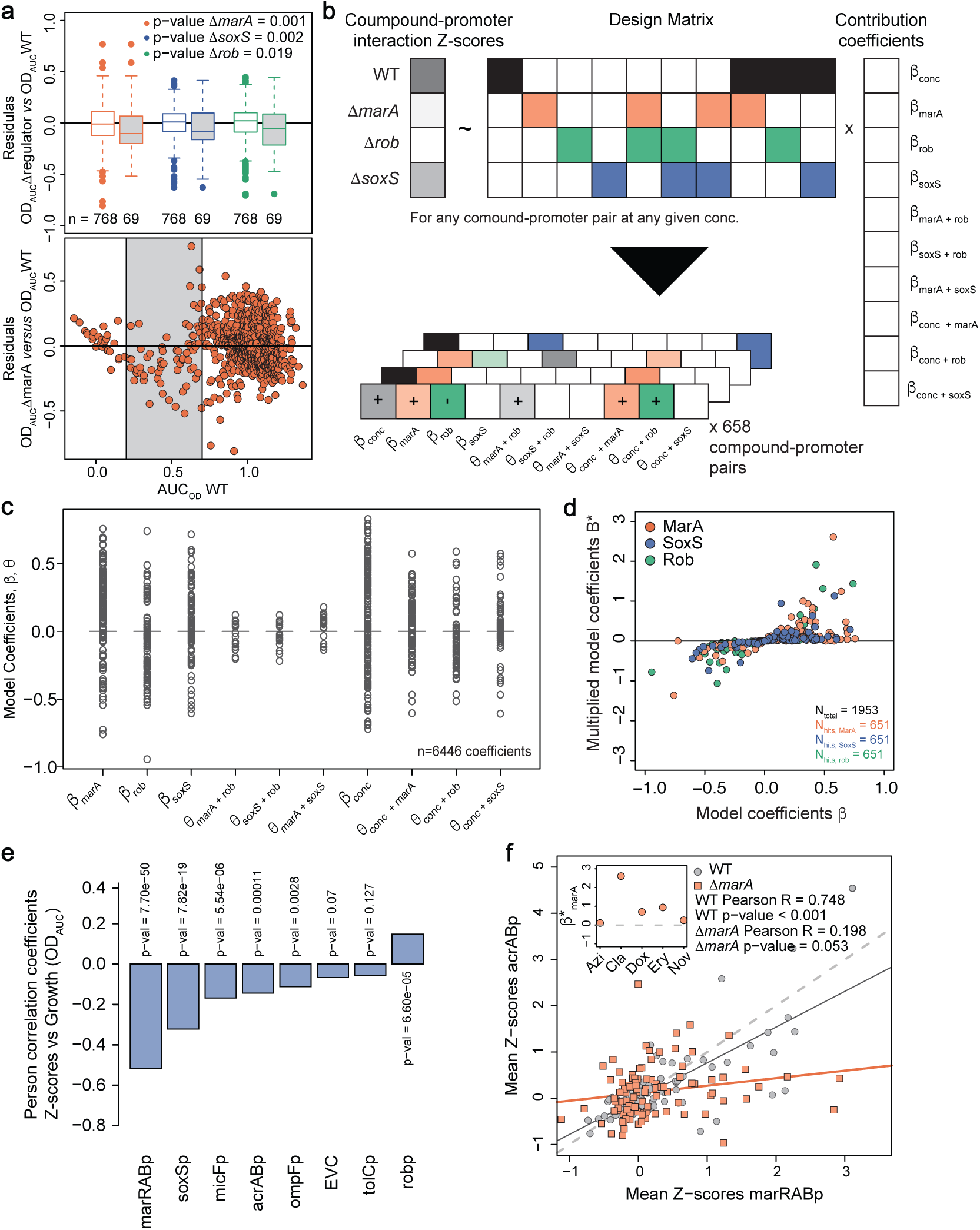
Assessing contributions of MarA, SoxS and Rob to compound- promoter interactions. **a)** Deletion of *marA*, *soxS* or *rob* sensitizes bacteria to several compounds at sub-inhibitory concentrations. Top: boxplots of the residuals of the lines-of-best-fit between growth (OD_AUC_) of the regulator mutants and wild-type for all tested pairs including water (EVC) across all 4 concentrations and 2 biological replicates. Negative residuals represent compounds concentrations to which the regulator mutant is more sensitive than the wild-type. White and gray boxes represent all residuals and a subset of compound concentrations with 0.2 < wild- type OD_AUC_ < 0.7, respectively. The number of data points is indicated below each box plot. Boxplots indicate 25^th^, 50^th^ and 75^th^ percentiles, and whiskers extend up to 1.5 times the interquartile range (IQR) from the 25^th^ and 75^th^ percentiles. p-value from a one-sided statistical t-test comparing full and subset residuals per mutant are shown. Bottom: Residuals of the lines-of-best-fit between growth (OD_AUC_) of βmarA and wild-type for all tested pairs including water (EVC) across all 4 concentrations and 2 biological replicates plotted against growth of the wild-type (containing EVC). Gray region corresponds to 0.2 < wild-type OD_AUC_ < 0.7. **b)** Schematic of the Lasso regression model to estimate regulator contributions to CPIs. Details described in Methods. **c)** Boxplots of contribution coefficients β and 8 grouped by name. Boxplots indicate 25^th^, 50^th^ and 75^th^ percentiles, and whiskers extend up to 1.5 times the interquartile range (IQR) from the 25^th^ and 75^th^ percentiles. Due to the nature of the data – very sharply zero-centered – 25^th^, 50^th^ and 75^th^ overlap. **d)** Scatterplot of multiplied coefficients B* versus model coefficients β of single regulator contributions, colored by regulator. **e)** Correlation of acrABp promoter activity with growth is lost upon *marA* deletion. Pearson correlation coefficients of Z-scores versus growth (OD_AUC_) for each individual promoter in ΔmarA. Correlation p-value (double sided t-test) shown above bars. **f)** acrABp and marRABp promoter activities are strongly correlated in a MarA-dependent manner. The mean Z-scores of all pairs including water (n=384) of acrABp plotted against marRABp. Correlation p-values (double sided t-test) and Pearson correlation coefficients are shown. Inlay shows β_marA_ to acrABp promoter activity in the wild-type for all CPIs involving acrABp.

**Extended data figure 4:**
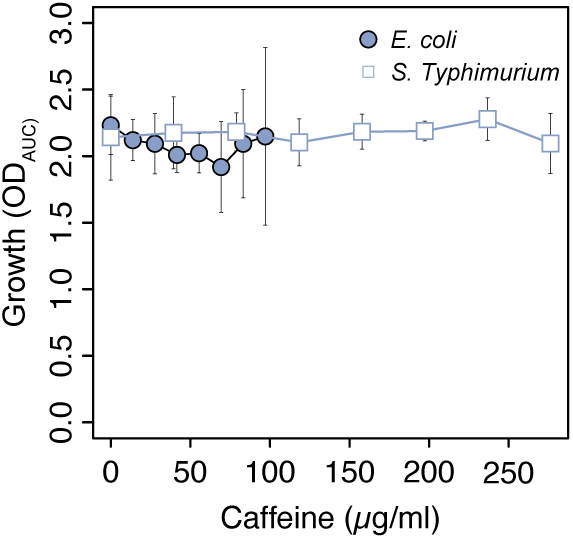
Caffeine does not inhibit growth of *E. coli* or *S.* Typhimurium. Caffeine MIC curves (growth vs caffeine concentration) of *E. coli* and *S.* Typhimurium wild-type strains. Mean growth (OD_AUC_, dots) and standard deviations (error bars) over 3 and 4 biological replicates for *E. coli* and *S.* Typhimurium, respectively.

## References

1. Nikaido, H. Molecular basis of bacterial outer membrane permeability revisited. Microbiol Mol Biol Rev 67, 593–656 (2003).

2. Du, D. et al. Multidrug efflux pumps: structure, function and regulation. Nat Rev Microbiol 16, 523–539 (2018).

3. Pagès, J.-M., James, C. E. & Winterhalter, M. The porin and the permeating antibiotic: a selective diffusion barrier in Gram-negative bacteria. Nat Rev Microbiol 6, 893–903 (2008).

4. Masi, M., Refregiers, M., Pos, K. M. & Pages, J. M. Mechanisms of envelope permeability and antibiotic influx and efflux in Gram-negative bacteria. Nat Microbiol 2, 17001 (2017).

5. Vergalli, J. et al. Porins and small-molecule translocation across the outer membrane of Gram-negative bacteria. Nat Rev Microbiol 18, 164–176 (2020).

6. Wang-Kan, X. et al. Lack of AcrB Efflux Function Confers Loss of Virulence on *Salmonella enterica* Serovar Typhimurium. mBio 8, e00968–17 (2017).

7. Pu, Y. et al. Enhanced Efflux Activity Facilitates Drug Tolerance in Dormant Bacterial Cells. Molecular Cell 62, 284–294 (2016).

8. El Meouche, I., Siu, Y. & Dunlop, M. J. Stochastic expression of a multiple antibiotic resistance activator confers transient resistance in single cells. Sci Rep 6, 19538 (2016).

9. El Meouche, I. & Dunlop, M. J. Heterogeneity in efflux pump expression predisposes antibiotic-resistant cells to mutation. Science 362, 686–690 (2018).

10. Maier, L. et al. Extensive impact of non-antibiotic drugs on human gut bacteria. Nature 555, 623–628 (2018).

11. Bioaccumulation of therapeutic drugs by human gut bacteria | Nature. https://www.nature.com/articles/s41586-021-03891-8.

12. Ricaurte, D. et al. High-throughput transcriptomics of 409 bacteria–drug pairs reveals drivers of gut microbiota perturbation. Nat Microbiol 9, 561–575 (2024).

13. Brochado, A. R. et al. Species-specific activity of antibacterial drug combinations. Nature 559, 259–263 (2018).

14. Cacace, E. et al. Systematic analysis of drug combinations against Gram-positive bacteria. Nat Microbiol 8, 2196–2212 (2023).

15. Chubiz, L. M. The Mar, Sox, and Rob Systems. EcoSal Plus eesp00102022 (2023) doi:10.1128/ecosalplus.esp-0010-2022.

16. Rosenberg, E. Y., Bertenthal, D., Nilles, M. L., Bertrand, K. P. & Nikaido, H. Bile salts and fatty acids induce the expression of Escherichia coli AcrAB multidrug efflux pump through their interaction with Rob regulatory protein. Mol Microbiol 48, 1609–19 (2003).

17. Nunoshiba, T., Hidalgo, E., Amabile Cuevas, C. F. & Demple, B. Two-stage control of an oxidative stress regulon: the Escherichia coli SoxR protein triggers redox-inducible expression of the soxS regulatory gene. J Bacteriol 174, 6054–60 (1992).

18. Cohen, S. P., Levy, S. B., Foulds, J. & Rosner, J. L. Salicylate induction of antibiotic resistance in Escherichia coli: activation of the mar operon and a mar-independent pathway. J Bacteriol 175, 7856–7862 (1993).

19. Barbosa, T. M. & Levy, S. B. Differential Expression of over 60 Chromosomal Genes in *Escherichia coli* by Constitutive Expression of MarA. J Bacteriol 182, 3467–3474 (2000).

20. Bennik, M. H. J., Pomposiello, P. J., Thorne, D. F. & Demple, B. Defining a *rob* Regulon in *Escherichia coli* by Using Transposon Mutagenesis. J Bacteriol 182, 3794–3801 (2000).

21. Pomposiello, P. J., Bennik, M. H. & Demple, B. Genome-wide transcriptional profiling of the Escherichia coli responses to superoxide stress and sodium salicylate. J Bacteriol 183, 3890–902 (2001).

22. Martin, R. G. & Rosner, J. L. Genomics of the marA/soxS/rob regulon of Escherichia coli: identification of directly activated promoters by application of molecular genetics and informatics to microarray data. Mol Microbiol 44, 1611–1624 (2002).

23. Sharma, P. et al. The multiple antibiotic resistance operon of enteric bacteria controls DNA repair and outer membrane integrity. Nat Commun 8, 1444 (2017).

24. Alekshun, M. N. & Levy, S. B. Alteration of the repressor activity of MarR, the negative regulator of the Escherichia coli marRAB locus, by multiple chemicals in vitro. J Bacteriol 181, 4669–72 (1999).

25. Rosner, J. L., Dangi, B., Gronenborn, A. M. & Martin, R. G. Posttranscriptional activation of the transcriptional activator Rob by dipyridyl in Escherichia coli. J Bacteriol 184, 1407– 16 (2002).

26. Alekshun, M. N., Levy, S. B., Mealy, T. R., Seaton, B. A. & Head, J. F. The crystal structure of MarR, a regulator of multiple antibiotic resistance, at 2.3 angstrom resolution. Nature Structural Biology 8, 710–714 (2001).

27. Kwon, H. J., Bennik, M. H., Demple, B. & Ellenberger, T. Crystal structure of the Escherichia coli Rob transcription factor in complex with DNA. Nat Struct Biol 7, 424–30 (2000).

28. Teelucksingh, T. et al. A genetic platform to investigate the functions of bacterial drug efflux pumps. Nat Chem Biol 18, 1399–1409 (2022).

29. Choi, U. & Lee, C.-R. Distinct Roles of Outer Membrane Porins in Antibiotic Resistance and Membrane Integrity in Escherichia coli. Front. Microbiol. 10, 953 (2019).

30. Zheng, M. & Lupoli, T. J. Counteracting antibiotic resistance enzymes and efflux pumps. Current Opinion in Microbiology 75, 102334 (2023).

31. Ramani, N., Hedeshian, M. & Freundlich, M. micF antisense RNA has a major role in osmoregulation of OmpF in Escherichia coli. J Bacteriol 176, 5005–10 (1994).

32. Goh, E.-B. et al. Transcriptional modulation of bacterial gene expression by subinhibitory concentrations of antibiotics. Proceedings of the National Academy of Sciences 99, 17025–17030 (2002).

33. Andersson, D. I. & Hughes, D. Microbiological effects of sublethal levels of antibiotics. Nat Rev Microbiol 12, 465–478 (2014).

34. Sambrook, J., Fritsch, E. F., Maniatis, T., Russell, D. W. & Green, M. R. Molecular Cloning: A Laboratory Manual. (Cold Spring Harbor Laboratory Press, Cold Spring Harbor, NY, 1989).

35. Rampersaud, A. & Inouye, M. Procaine, a local anesthetic, signals through the EnvZ receptor to change the DNA binding affinity of the transcriptional activator protein OmpR. J Bacteriol 173, 6882–6888 (1991).

36. Schneiders, T. & Levy, S. B. MarA-mediated Transcriptional Repression of the rob Promoter. Journal of Biological Chemistry 281, 10049–10055 (2006).

37. Chubiz, L. M., Glekas, G. D. & Rao, C. V. Transcriptional cross talk within the mar-sox-rob regulon in Escherichia coli is limited to the rob and marRAB operons. J Bacteriol 194, 4867–75 (2012).

38. Michán, C., Manchado, M. & Pueyo, C. SoxRS Down-Regulation of *rob* Transcription. J Bacteriol 184, 4733–4738 (2002).

39. Hächler, H., Cohen, S. P. & Levy, S. B. marA, a regulated locus which controls expression of chromosomal multiple antibiotic resistance in Escherichia coli. J Bacteriol 173, 5532– 5538 (1991).

40. Hao, Z. et al. The multiple antibiotic resistance regulator MarR is a copper sensor in Escherichia coli. Nat Chem Biol 10, 21–28 (2014).

41. Chubiz, L. M. & Rao, C. V. Role of the mar-sox-rob regulon in regulating outer membrane porin expression. J Bacteriol 193, 2252–60 (2011).

42. Okusu, H., Ma, D. & Nikaido, H. AcrAB efflux pump plays a major role in the antibiotic resistance phenotype of Escherichia coli multiple-antibiotic-resistance (Mar) mutants. J Bacteriol 178, 306–308 (1996).

43. Cohen, S. P., McMurry, L. M. & Levy, S. B. marA locus causes decreased expression of OmpF porin in multiple-antibiotic-resistant (Mar) mutants of Escherichia coli. J Bacteriol 170, 5416–22 (1988).

44. Bien, J., Taylor, J. & Tibshirani, R. A lasso for hierarchical interactions. Ann. Statist. 41, (2013).

45. Tibshirani, R. Regression Shrinkage and Selection Via the Lasso. Journal of the Royal Statistical Society Series B: Statistical Methodology 58, 267–288 (1996).

46. Ruiz, C. & Levy, S. B. Many Chromosomal Genes Modulate MarA-Mediated Multidrug Resistance in *Escherichia coli*. Antimicrob Agents Chemother 54, 2125–2134 (2010).

47. Ruiz, C. & Levy, S. B. Regulation of acrAB expression by cellular metabolites in Escherichia coli. Journal of Antimicrobial Chemotherapy 69, 390–399 (2014).

48. Piddock, L. J. V. Fluoroquinolone resistance in Salmonella serovars isolated from humans and food animals1. FEMS Microbiology Reviews 26, 3–16 (2002).

49. Maher, C. & Hassan, K. A. The Gram-negative permeability barrier: tipping the balance of the in and the out. mBio 14, e01205–23 (2023).

50. Li, X.-Z., Plésiat, P. & Nikaido, H. The Challenge of Efflux-Mediated Antibiotic Resistance in Gram-Negative Bacteria. Clinical Microbiology Reviews 28, 337–418 (2015).

51. Ricci, V., Kaur, J., Stone, J. & Piddock, L. J. V. Antibiotics do not induce expression of acrAB directly but via a RamA-dependent pathway. Antimicrob Agents Chemother 67, e0062023 (2023).

52. Martin, R. G., Gillette, W. K., Rhee, S. & Rosner, J. L. Structural requirements for marbox function in transcriptional activation of *mar* / *sox* / *rob* regulon promoters in *Escherichia coli* : sequence, orientation and spatial relationship to the core promoter. Molecular Microbiology 34, 431–441 (1999).

## References

1. Uehara, T., Parzych, K. R., Dinh, T. & Bernhardt, T. G. Daughter cell separation is controlled by cytokinetic ring-activated cell wall hydrolysis. EMBO J. 29, 1412–1422 (2010).

2. Baba, T. et al. Construction of Escherichia coli K-12 in-frame, single-gene knockout mutants: the Keio collection. Mol Syst Biol 2, 2006 0008 (2006).

3. Porwollik, S. et al. Defined Single-Gene and Multi-Gene Deletion Mutant Collections in Salmonella enterica sv Typhimurium. PLOS ONE 9, e99820 (2014).

4. Datsenko, K. A. & Wanner, B. L. One-step inactivation of chromosomal genes in Escherichia coli K-12 using PCR products. Proc. Natl. Acad. Sci. U. S. A. 97, 6640–6645 (2000).

5. Huber, P. J. Robust Statistics. (Wiley, 1981). doi:10.1002/0471725250.

6. Benjamini, Y. & Hochberg, Y. Controlling the False Discovery Rate: A Practical and Powerful Approach to Multiple Testing. J. R. Stat. Soc. Ser. B Stat. Methodol. 57, 289– 300 (1995).

7. Livak, K. J. & Schmittgen, T. D. Analysis of Relative Gene Expression Data Using Real- Time Quantitative PCR and the 2−ΔΔCT Method. Methods 25, 402–408 (2001).

8. Tibshirani, R. Regression Shrinkage and Selection Via the Lasso. J. R. Stat. Soc. Ser. B Stat. Methodol. 58, 267–288 (1996).

9. Bien, J., Taylor, J. & Tibshirani, R. A lasso for hierarchical interactions. (2012) doi:10.48550/ARXIV.1205.5050.

10. Fuchs, M. et al. A network of small RNAS regulates sporulation initiation in *Clostridioides difficile*. EMBO J. 42, e112858 (2023).

11. Schneider, C. A., Rasband, W. S. & Eliceiri, K. W. NIH Image to ImageJ: 25 years of image analysis. Nat. Methods 9, 671–675 (2012).

12. Schindelin, J., et al. Fiji: an open-source platform for biological-image analysis. Nat. Methods 9, 676–682 (2012).

